# Modeling of Ion and Water Transport in the Biological Nanopore ClyA

**DOI:** 10.1101/2020.01.08.897819

**Authors:** Kherim Willems, Dino Ruić, Florian Lucas, Ujjal Barman, Johan Hofkens, Giovanni Maglia, Pol Van Dorpe

## Abstract

In recent years, the protein nanopore cytolysin A (ClyA) has become a valuable tool for the detection, characterization and quantification of biomarkers, proteins and nucleic acids at the single-molecule level. Despite this extensive experimental utilization, a comprehensive computational study of ion and water transport through ClyA is currently lacking. Such a study yields a wealth of information on the electrolytic conditions inside the pore and on the scale the electrophoretic forces that drive molecular transport. To this end we have built a computationally efficient continuum model of ClyA which, together with an extended version of Poison-Nernst-Planck-Navier-Stokes (ePNP-NS) equations, faithfully reproduces its ionic conductance over a wide range of salt concentrations. These ePNP-NS equations aim to tackle the shortcomings of the traditional PNP-NS models by self-consistently taking into account the influence of both the ionic strength and the nanoscopic scale of the pore on all relevant electrolyte properties. In this study, we give both a detailed description of our ePNP-NS model and apply it to the ClyA nanopore. This enabled us to gain a deeper insight into the influence of ionic strength and applied voltage on the ionic conductance through ClyA and a plethora of quantities difficult to assess experimentally. The latter includes the cation and anion concentrations inside the pore, the shape of the electrostatic potential landscape and the magnitude of the electro-osmotic flow. Our work shows that continuum models of biological nanopores—if the appropriate corrections are applied—can make both qualitatively and quantitatively meaningful predictions that could be valuable tool to aid in both the design and interpretation of nanopore experiments.

## Introduction

The transport of ions and molecules through nanoscale geometries is a field of intense study that uses both experimental, theoretical and computational methods. ^1–6^ One of the primary driving forces behind this research is the development of nanopores as label-free, stochastic sensors at the ultimate analytical limit (*i.e.*, single molecule).^7–10^ Such detectors have applications ranging from the analysis of biopolymers such as DNA^11–18^ or proteins,^19–23^ to the detection and quantification of biomarkers,^24–30^ to the fundamental study of chemical or enzymatic reactions at the single molecular level. ^22,31–34^

Nanopores are typically operated in the resistive-pulse mode, where the fluctuations of their ionic conductance are monitored over time.^7–9,35^ Experimentally, this is achieved by placing the nanopore between two electrolyte compartments and applying a constant DC (or AC) voltage across them. Due to the high resistance of the nanopore, virtually the full potential change occurs within (and around) the pore, resulting in a strong electric field (10^6^–10^7^ mV · nm^−1^) that electrophoretically drives ions and water molecules through it.^36–39^ Hence, analyte molecules such as DNA or proteins are driven towards, and often *through*, the nanopore by a combination of Coulombic (electrophoretic) and hydrodynamic (electro-osmotic) forces.^36,40–42^ If successful, a translocation event is observed as a temporal fluctuation in the ionic conductance of the pore that serves as a unique molecular ‘fingerprint’ with which the molecule can be identified.^43^ Because the frequency, magnitude, duration and even noise levels of these events depend on the properties of both the analyte molecule and the nanopore itself, they are notoriously difficult to interpret unambiguously without a full understanding of the nanofluidic phenomena that underlie them.

The computational approaches most widely used to study nanofluidic transport in ion channels or biological nanopores comprise *discrete* methods such as molecular dynamics (MD)^44–51^ and Brownian dynamics (BD),^52–58^ and *mean-field* (continuum) methods based on solving the Poisson-Boltzmann (PB) equations^59,60^ and Poisson-Nernst-Planck (PNP) equations. ^61–63^ The latter two can be coupled with the Navier-Stokes (NS) equation to include electro-osmotically or pressure driven fluid flow. ^58,64^ Due to their explicit atomic or particle nature, MD and BD simulations are considered to yield the most accurate results. However, the large computational cost of simulating a complete biological nanopore system (100K–1M atoms) for hundreds of nanoseconds still necessitates the use of supercomputers.^46,48^ The PNP(-NS) equations, on the other hand, are of particular interest due to their low computational demands and analytical tractability. In a continuum approach, the simulated system is subdivided in several ‘structureless’ domains, the behavior of which is parameterized by material properties such as relative permittivity, diffusion coefficient, electrophoretic mobility, viscosity and density. Because these properties can only emerge from the collective behavior or interactions between small groups of atoms (*i.e.*, the mean-field approximation), great care must be taken when using them to compute fluxes and fields at the nanoscale, where computational elements may only contain a few molecules. ^65,66^ Nevertheless, the PNP equations have been used extensively for the simulation of ion channels, ^53,67,68^ biological nanopores^58,63,69,70^ and their solid-state counterparts^39,71–73^—often with excellent qualitative, if not quantitative results.^3,4,6^

To remedy the shortcomings of PNP and NS theory, a number of modifications have been proposed over the years. These include, among others, 1) steric ion-ion interactions, 2) the effect of protein-ion/water interactions on their motility (*i.e.*, diffusivity and electrophoretic mobility), 3) the concentration dependencies of ion motility, and solvent relative permittivity, viscosity and density.

The steric ion-ion interactions can be accounted for by computing the excess in chemical potential 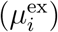 resulting from the finite size of the ions.^61,74,75^ Gillespie *et al.* combined PNP and density functional theory—where 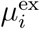 was split up in ideal, hard-sphere and electrostatic components—to successfully predict the selectivity and current of ion channels. ^62^ In another approach, Kilić *et al.* derived a set of modified PNP equations based on the free energy functional of the Borukhov’s modified PB model ^76^ and observed significantly more realistic concentrations for high surface potentials compared to the classical PNP equations. ^77^ To allow for non-identical ion sizes and more than two ion species, this model was later extended by Lu *et al.*, who used it to probe the effect of finite ion size on the rate coefficients of enzymes. ^78^

The interaction of ions or small molecules such as water with the heavy atoms of proteins or DNA results in a strong reduction of their motility, as observed in MD simulations.^79,80^ Since these effects happen only at distances ≤1 nm, they can usually be neglected for macroscopic simulations. However, in small nanopores (≤ 10 nm radius), they comprise a significant fraction of the total nanopore radius and hence must be taken into account.^54,58,63,81^ In continuum simulations, this can be achieved with the use pf positional-dependent ion diffusion coefficients. An example implementation is the ‘soft-repulsion PNP’ developed by Simakov and Kurnikova, ^63,70^ who used it to predict the ionic conductance of the *α*-hemolysin nanopore. Similar reductions in ion diffusion coefficients have been proposed to improve PNP theory’s estimations of the ionic conductance of ion channels. ^67,68^ The motility of water molecules is expressed by the NS equations as the fluid’s viscosity. Hence, as also observed in MD simulations for water molecules near proteins^80^ and confined in hydrophilic nanopores,^82–84^ the water-solid interaction leads to a viscosity several times higher compared to the bulk values. Note that this is valid for hydrophilic interfaces only, as the lack of interaction with hydrophobic interfaces, such as carbon nanotubes, leads to a lower viscosity.^85^

It is well known that the self-diffusion coefficient 𝒟_*i*_ and electrophoretic mobility *µ*_*i*_ of an ion *i* depends on the local concentrations of all the ions in the electrolyte. ^86^ Their values typically decrease with increasing salt concentration, and should not be treated as constants. Moreover, even though the well-known Nernst-Einstein (NE) relation *µ*_*i*_ = 𝒟_*i*_*/k*_B_*T* is strictly speaking only valid at infinite dilution and a good approximation at low concentrations (<10 mM), it significantly overestimates the ionic mobility at higher salt concentrations. ^86–89^ In an empirical approach, Baldessari and Santiago formulated an ionic-strength dependency of the ionic mobility based on the activity coefficient of the salt^60^ and showed excellent correspondence between the experimental and simulated ionic conductance of long nanochannels over a wide concentration range.^90^ Alternatively, Burger *et al.* used a microscopic lattice-based model to derive a set of PNP equations with non-linear, ion density-dependent mobilities and diffusion coefficients that provided significantly more realistic results for ion channels.^91^ Note that other electrolyte properties, such as its viscosity, ^92^ density^92^ and relative permittivity, ^93^ also significantly affect the ion and water flux. To better compute the charge flux in ion channels, Chen derived a new PNP framework^94^ that includes water-ion interactions in the form of a concentration-dependent relative permittivity and an additional ion-water interaction energy term.

To the best of our knowledge, no attempt has been made to consolidate all of the corrections discussed above into a single framework. Hence, we propose an extended set of PNP-NS (ePNP-NS) equations, which improves the predictive power of the PNP-NS equations at the nanoscale and beyond infinite dilution. Our ePNP-NS framework takes into account the finite size of the ions using a size-modified PNP theory,^78^ and implements spatial-dependencies for the solvent viscosity,^80,84^ the ion diffusion coefficients and their mobilities.^54,79^ It also includes self-consistent concentration-dependent properties (based on empirical fits to experimental data) for both all ions in terms of diffusion coefficients and mobilities,^60,87^ and the solvent in terms of density, viscosity^92^ and relative permittivity^93^. To validate our new framework, we applied it directly to a 2D-axisymmetric model of Cytolysin A (ClyA), a large protein nanopore that typically contains 12 subunits^95^ or more^96^ and has been extensively used in experimental studies of both proteins^27,30,96–100^ and DNA.^17,101^. This allowed us to gauge the qualitative and quantitative performance of the ePNP-NS equations and simultaneously elucidate previously unaccessible details about the environment inside the pore.

The remainder of this paper is organized as follows. In *Mathematical model* we describe the equations governing our ePNP-NS framework and detail the construction of the 2D-axisymmetric ClyA model. Next, in *Results and Discussion*, we validate our model by direct comparison of simulated ionic conductance with experimentally measured values. We then proceed to characterize the influence of the bulk ionic strength and the applied bias voltage on cation and anion concentrations inside the pore, the electrostatic potential distribution and magnitude of the electro-osmotic flow. Finally, we touch upon our key finding and their impact in *Conclusions* and describe our protocols in more detail in *Materials and Methods*.

## Mathematical model

The use of *continuum* or *mean-field* representations for both the nanopore and the electrolyte enables us to efficiently compute the steady-state ion and water fluxes under almost any condition. The dynamic behavior of our complete system is described by the coupled Poisson, Nernst-Planck and Navier-Stokes (PNP-NS) equations, a well-known set of partial differential equations that describe the electrostatic field, the total ionic flux and the fluid flow, respectively. ^61,64,71^ In this section we will describe all the components of the simulation, *i.e.*, the 2D-axisymmetric nanopore geometry and the system of governing equations.

### Model geometry

#### 2D-axisymmetric model of ClyA

ClyA is a relatively large protein nanopore that self-assembles on lipid bilayers to form 14 nm long hydrophilic channels. The interior of the pore can be divided into roughly two cylindrical compartments (Fig. 1a): the *cis lumen* (≈6 nm diameter, ≈10 nm height), and the *trans* constriction (≈3.3 nm diameter, ≈4 nm height). Because ClyA consists of 12 identical subunits (Fig. 1b), it exhibits a high degree of radial symmetry, a geometrical feature that can be exploited to obtain meaningful results at a much lower computational cost. ^58,64,71^ However, this requires the reduction of the full 3D atomic structure and charge distribution to a realistic 2D-axisymmetric model. To this end, we constructed a full-atom homology model of ClyA-AS type I—a dodecameric variant of the wild type ClyA from *S. Typhii* artificially evolved for improved stability^96^—and equilibrated it at 298.15 K for 30 ns in an explicit solvent with harmonic contraints on the protein backbone atoms (see Materials and Methods for details). From the final 5 ns of this trajectory we extracted 50 sets of atomic coordinates for ClyA (*i.e.*, every 100 ps), in order to adequately represent the conformational diversity of the side chains. For each of these structures, we computed a 2D atomic density^104^ and charge^46^ map (see Materials and Methods for details), and the final maps were obtained through averaging. The geometry of the nanopore was then defined as the 25 % contour line of the density map, which closely follows the outline of a 30° ‘wedge’ of the full atom structure (Fig. 1c). The equilibrium charge map (Fig. 1d) was loaded directly into our solver as a linear interpolation function 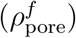 and applied across all computational domains.

**Figure 1.**
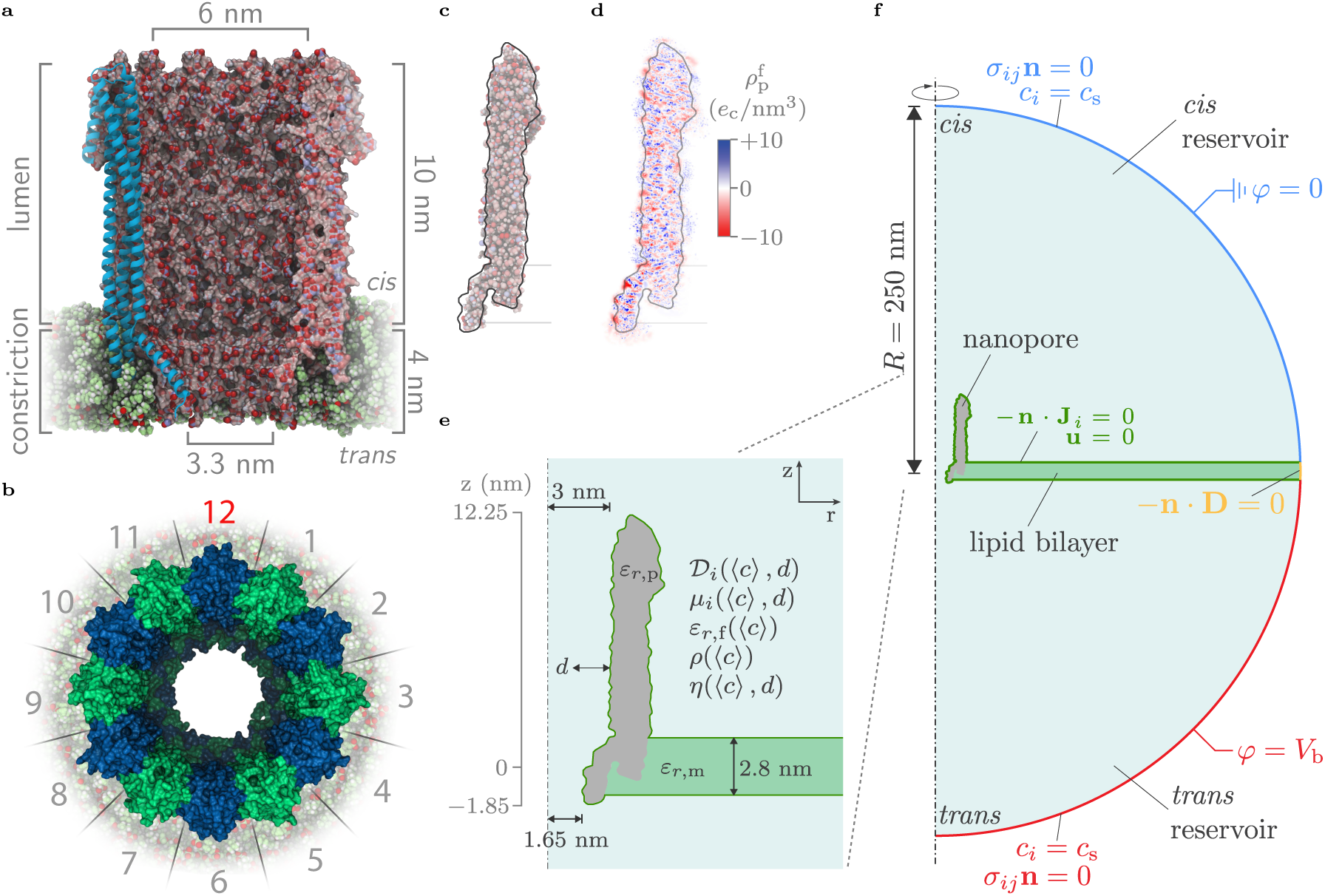
All-atom and 2D-axisymmetric models of ClyA. (a) Axial cross-sectional and (b) top views of the dodecameric nanopore ClyA-AS,^96^ derived through homology modelling from the *E. coli* Cytolysin A crystal structure (PDBID: 2WCD^95^). Figures were rendered with VMD.^102,103^ (c) The 2D-axisymmetric geometry was derived directly from the all-atom model by computing the average inner and outer radii along the longitudinal axis of the pore, and hence closely follows the outline of a 30° wedge out of the homology model. (d) The fixed space charge density 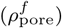 map of ClyA-AS, obtained by Gaussian projection of each atom’s partial charge onto a 2D plane (see methods for details). (e+f) The 2D-axisymmetric simulation geometry of ClyA (grey) embedded in a lipid bilayer (green) and surrounded by a spherical water reservoir (blue). Note that all electrolyte parameters depend on the local average ion concentration 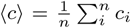 and that some are also influenced by the distance from the nanopore wall *d*.

#### Global geometry

The complete system (Figs. 1e and 1f) consists of a large hemispherical electrolyte reservoir (*R* = 250 nm), split through the middle into a *cis* and a *trans* compartment by a lipid bilayer (*h* = 2.8 nm), which contains the nanopore at its center. Both the bilayer and the nanopore are represented by dielectric blocks (see Tab. 1 for parameters) that are impermeable to ions and water.

**Table 1.** Summary of the parameters and fitting equations used in the ePNP-NS equations.

| Property | Parameter<sup>a</sup> | Infinite dilution value<sup>b</sup>/Fitting function<sup>c</sup> | Reference |
| --- | --- | --- | --- |
| Relative permittivity | $\varepsilon_{r,p}$ | 20 | 104 |
| | $\varepsilon_{r,m}$ | 3.2 | 105 |
| | $\varepsilon_{r,f}^0$ | 78.15 | 93 |
| | $\varepsilon_w(\bar{c})$ | $1 - \left(1 - \frac{P_1}{P_0}\right) L\left(\frac{3P_2}{P_0 - P_1}\bar{c}\right)$ | 93 |
| Ion self-diffusion coefficient | $\mathcal{D}_{Na^+}^0$ | $1.334 \times 10^{-9} \text{ m}^2 \cdot \text{s}^{-1}$ | 87 |
| | $\mathcal{D}_{Cl^-}^0$ | $2.032 \times 10^{-9} \text{ m}^2 \cdot \text{s}^{-1}$ | 87 |
| | $\mathcal{D}_i^c(\bar{c})$ | $(1 + P_1\bar{c}^{0.5} + P_2\bar{c} + P_3\bar{c}^{1.5} + P_4\bar{c}^2)^{-1}$ | This work |
| | $\mathcal{D}_i^w(\bar{d})$ | $1 - \exp(-P_1(\bar{d} + P_2))$ | 63,79 |
| Ion electrophoretic mobility | $\mu_{Na^+}^0$ | $5.192 \times 10^{-4} \text{ m}^2 \cdot \text{s}^{-1} \cdot \text{V}^{-1}$ | 106 |
| | $\mu_{Cl^-}^0$ | $7.909 \times 10^{-4} \text{ m}^2 \cdot \text{s}^{-1} \cdot \text{V}^{-1}$ | 106 |
| | $\mu_i^c(\bar{c})$ | $(1 + P_1\bar{c}^{0.5} + P_2\bar{c} + P_3\bar{c}^{1.5} + P_4\bar{c}^2)^{-1}$ | This work |
| | $\mu_i^w(\bar{d})$ | $1 - \exp(-P_1(\bar{d} + P_2))$ | 63,79 |
| Ion transport number | $t_{Na^+}^0$ | 0.396 | 106 |
| | $t_{Na^+}(\bar{c})$ | $(1 + P_1\bar{c}^{0.5} + P_2\bar{c} + P_3\bar{c}^{1.5} + P_4\bar{c}^2)^{-1}$ | This work |
| Dynamic viscosity | $\eta^0$ | $8.904 \text{ Pa} \cdot \text{s}$ | 92 |
| | $\eta^c(\bar{c})$ | $1 + P_1\bar{c}^{0.5} + P_2\bar{c} + P_3\bar{c}^2 + P_4\bar{c}^{3.5}$ | This work |
| | $\eta^w(\bar{d})$ | $1 + \exp(-P_1(\bar{d} - P_2))$ | 80 |
| Fluid density | $\varrho^0$ | $997 \text{ kg} \cdot \text{m}^{-3}$ | 92 |
| | $\varrho(\bar{c})$ | $1 + P_1\bar{c} + P_2\bar{c}^2$ | This work |

<sup>a</sup>Dependencies on either $\bar{c} = \langle c \rangle / 1 \text{ M}$ (dimensionless average ion concentration) and $\bar{d} = d / 1 \text{ nm}$ (dimensionless distance from the nanopore wall); <sup>b</sup>Values at infinite dilution for a system temperature of 291.15 K; <sup>c</sup>These functions are empirical and hence have no physical meaning. The values of the fitting parameters $P_x$ of each property can be found in Tab. S2 and graphs of the fits in Fig. S1.

### Governing equations

In an attempt to improve upon the quantitative accuracy of the PNP-NS equations for nanopore simulations, we developed an *extended* version of these equations (ePNP-NS) and implemented it in the commercial finite element solver COMSOL Multiphysics (v5.4, www.comsol.com). Our ePNP-NS equations self-consistently take into account 1) the finite size of the ions,^76,78^ 2) the reduction of ion and water motility close to the nanopore walls,^54,58,79,80,83^ and 3) the concentration dependency of ion diffusion coefficients and electrophoretic mobilities, as well as electrolyte viscosity, density and relative permittivity. ^87,92,93^ Most of these corrections make use of using empirical functions that were fitted to experimental data (Tabs. 1 and S2).

#### Electrostatic field

As mentioned above, the electrostatic potential is evaluated using Poisson’s equation

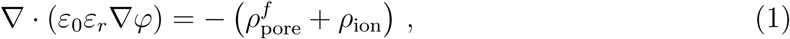

with *φ* the electric potential, *ε*_0_ the vacuum permittivity (8.854 19×10^−12^ F · m^−1^) and *ε*_*r*_ local relative permittivity (Tab. S2).

The pore’s *fixed* charge distribution, 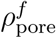, (see Eq. 18) was derived directly from the full atom model of ClyA-AS. The *ionic* charge density in the fluid is given by

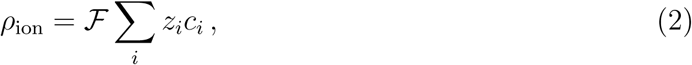

with ℱ Faraday’s constant (96 485.33 C · mol^−1^), and *c*_*i*_ the ion concentration and *z*_*i*_ ion charge number of ion *i*.

To account for the concentration dependence of the electrolyte’s relative permittivity, we used the expression

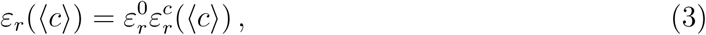

with 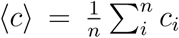 the average ion concentration, 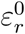 the relative permittivity at infinite dilution and 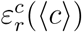 a concentration dependent empirical function parameterized with experimental data.

#### Ionic flux

The total ionic flux ***J***_*i*_ of each ion *i* is given by the size-modified Nernst-Planck equation, ^78^ and can be expressed as the sum of diffusive, electrophoretic, convective and steric fluxes

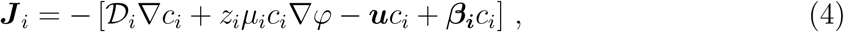

where

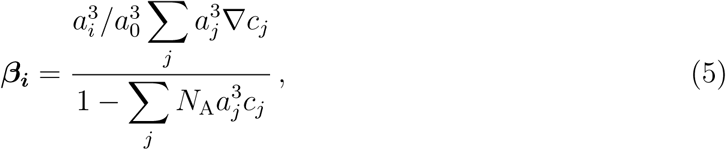

and at steady state

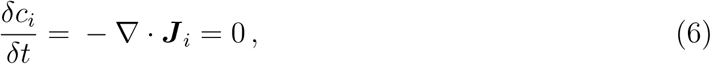

with 𝒟_*i*_ the ion diffusion coefficient, *c*_*i*_ the ion concentration, *z*_*i*_ the ion charge number, *µ*_*i*_ the electrophoretic mobility of ion *i. φ* is the electrostatic potential, ***u*** the fluid velocity and *N*_A_ Avogadro’s constant (6.022×10^23^ mol^−1^). *a*_*i*_ and *a*_0_ are *steric* cubic diameters of respectively ions and water molecules. Because currently there are no experimentally verified values available for *a*_*i*_ and *a*_0_, we set them to 0.5 nm (max. 13.3 M) and 0.311 nm (max. 55.2 M).^74^

The reduction of the ionic motility at increasing salt concentrations and in proximity to the nanopore walls was implemented self-consistently by replacing 𝒟_*i*_ and *µ*_*i*_ with the expressions

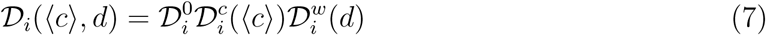

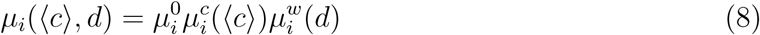

where 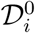 and 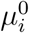 represent the values at infinite dilution. The concentration dependent factors 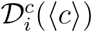 and 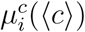 are empirical functions fitted to experimental data (between 0 to 5 M NaCl) of respectively the ion self-diffusion coefficients^87^ and the electrophoretic mobilities.^106–109^ Likewise, the factors 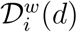 and 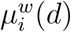 are empirical functions that introduce a spatial dependency on the distance from the nanopore wall *d*, and were parameterized by fitting to molecular dynamics data.^54,63,79^

Based on the observation that the diffusivity of nanometer- to micrometer-sized particles reduces significantly when confined in pores and slits of comparable dimensions, ^42,110–113^ Simakov *et al.*^63^ and Pederson et al.^58^ reduced the ion motilities inside the pore as a function of the ratio between the ion and the nanopore radii. We chose not to include this correction into our model, as extrapolating its applicability for ions with a hydrodynamic radii comparable to size of the solvent molecules is questionable.^111,114^

#### Fluid flow

The fluid flow and pressure are given by the Navier-Stokes equations for incompressible fluids^115^

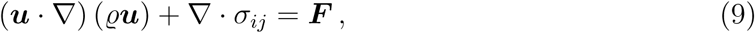

where

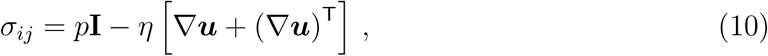

together with the continuity equations for the fluid density

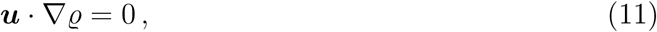

and the fluid velocity

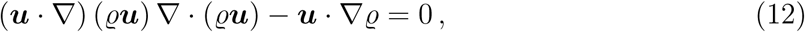

with ***u*** the fluid velocity, *ϱ* the fluid density, *σ*_*ij*_ the hydrodynamic stress tensor, *η* the viscosity and *p* the pressure. The external body force density ***F*** that acts on the fluid is given by

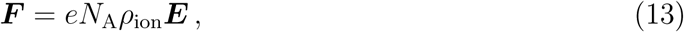

with ***E*** = −∇*φ* the electric field vector.

As with the previous equations, we introduced a concentration dependency and wall distance dependencies for *η* and a concentration dependency for the *ϱ* by replacing their constant values by

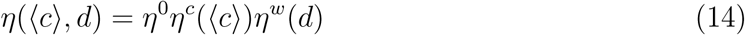

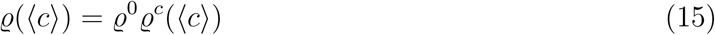

where *η*^0^ and *ϱ*^0^ are the values at infinite dilution (*i.e.*, pure water). The empirical functions *η*^*c*^(⟨*c*⟩), *ϱ*^*c*^(⟨*c*⟩) and *η*^*w*^(*d*) were parameterized *via* fitting to experimental^92^ and molecular dynamics^80^ data obtained from literature.

### Boundary conditions

The reservoir boundaries were set up, with Dirichlet conditions, to act as electrodes: the *cis* side was grounded (*φ* = 0) and a fixed, but changeable, bias potential was applied along the *trans* edge (*φ* = *V*_b_). To simulate the presence of an endless reservoir, the ion concentration at both external boundaries were fixed to the bulk salt concentration (*c*_*i*_ = *c*_s_) and the unconstrained flow in and out of the computational domain was enabled by means of a ‘no normal stress’ condition (*σ*_*ij*_***n*** = 0). The boundary conditions on the edges of the reservoir shared with the nanopore and bilayer were set to no-flux (−***n*** ·***J***_*i*_ = 0) and no-slip (***u*** = 0), preventing the flux of ions through them and mimicking a sticky hydrophilic surface, respectively. Finally, a Neumann boundary condition was applied at the bilayer’s external boundary (−***n*** · (*ε*_0_*ε*_*r*_∇*φ*) = 0).

### Note on the concentration dependencies

All concentration dependent parameters use the local ionic strength rather than their individual ion concentrations. Though valid for *electroneutral* bulk solutions, this approximation no longer holds inside the electrical double layer (*i.e.*, near charged surfaces or inside small nanopores), where local electroneutrality is violated. The main reasons for nevertheless making this simplification are the lack of non-bulk experimental data and the absence of a tractable analytical model. Furthermore, we will see that the current implementation of our concentration dependent functions will lead to an excellent agreement with the experimental data in all but the most extreme cases, justifying our choice *a posteriori*.

### PNP-NS vs. ePNP-NS

The ePNP-NS equations revert into the regular PNP-NS equations disabling the steric flux (***β*** = 0) and by setting all concentration and wall distance functions to unity (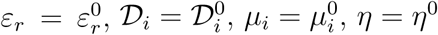 and *ϱ* = *ϱ*^0^).

## Results and Discussion

The current–voltage (IV) relationships of many nanopores, ClyA included, often deviate significantly from Ohm’s law. This is because the ionic flux arises from a complex interplay between the pore’s geometry (*e.g.*, size, shape and charge distribution), the properties of the surrounding electrolyte (*e.g.*, salt concentration viscosity and relative permittivity) and the externally applied conditions (*e.g.*, bias voltage, temperature and pressure). The ability of a computational model to quantitatively predict the ionic current of a nanopore over a wide range of bias voltages and salt concentrations strongly indicates that it captures the essential physics governing the nanofluidic transport. Hence, to validate our model, we experimentally measured the single channel ionic conductance of ClyA at a wide range of experimentally relevant salt concentrations and bias voltages. We compared these experimental data with the simulated ionic transport properties in terms of current, conductance, rectification and ion selectivity, of both the classical PNP-NS and the newly developed ePNP-NS equations. After validation, we proceed with describing the influence of the bias voltage (*V*_b_) and bulk salt concentration (*c*_s_) on the local ion and charge distribution and inside the pore. This is followed by a characterization of the electrostatic potential and the electrostatic energy landscape within ClyA for both cations and anions. We will conclude this section by discussing the properties of the electro-osmotic flow.

### Transport of Ions Through ClyA

#### Ionic current and conductance

The ability of our model to reproduce the ionic current of a biological nanopore over a wide range of experimentally relevant conditions (between *V*_b_ = −150 to +150 mV and for *c*_s_ = 0.05, 0.15, 0.5, 1 and 3 M NaCl) can be seen in Fig. 2a. Here, we compare IV relationships of ClyA-AS as measured experimentally (‘expt.’), simulated using our 2D-axisymmetric model (‘PNP-NS’ and ‘ePNP-NS’) and naively analytically estimated (‘bulk’) using a resistor model of the pore^96,116^ (Eqs. S18 and S19). Whereas the classical PNP-NS equations consistently overestimated the ionic current, particularly at high salt concentrations, the predictions of the ePNP-NS equations corresponded closely to the measured values, *especially* at high ionic strengths (*c*_s_ > 0.5 M). The inability of the classical PNP-NS equations to correctly estimate the current is expected however, as in this regime the model parameters (*e.g.*, diffusivity, mobility and viscosity, …) begin to deviate significantly from their ‘infinite dilution’ values (see Fig. S1). At lower salt concentrations the ePNP-NS equations tended to minorly overestimate the ionic current, but the discrepancies were much smaller than those observed for PNP-NS. Finally, the bulk model managed to capture the currents suprisingly well at high salt concentrations and positive bias voltages, but faltered in the negative voltage regime.

**Figure 2.**
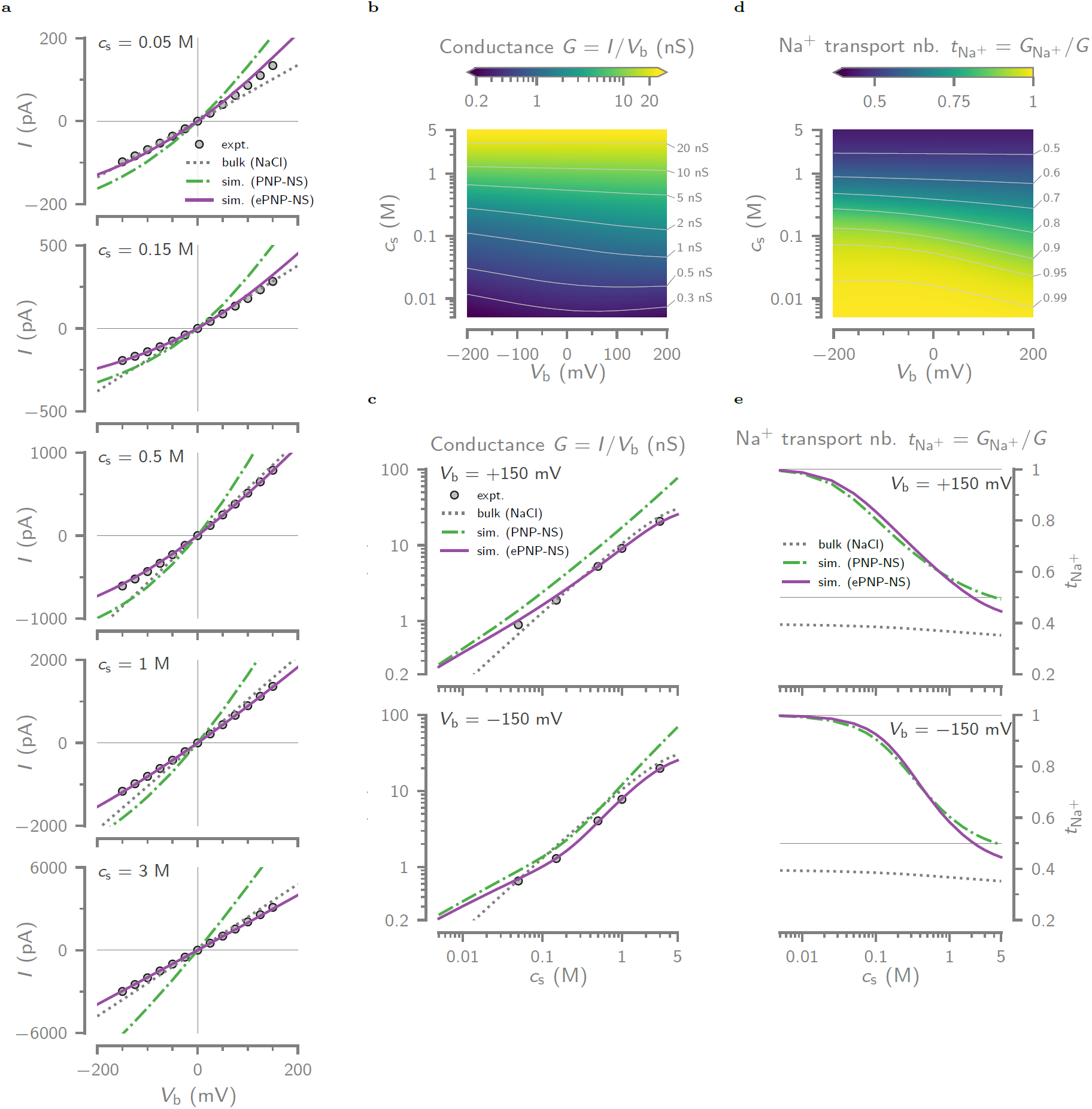
Measured and simulated ionic conductance and cation selectivity of single ClyA nanopores. (a) Comparison between the experimentally (expt.) measured, bulk pore model (bulk) and the simulated (PNP-NS and ePNP-NS) current-voltage (IV) curves of ClyA-AS at 25 ±1 °C between *V*_b_ = − 200 to +200 mV, and for *c*_s_ = 0.05, 0.15, 0.5, 1 and 3 M NaCl. The bulk current was calculated by Eq. S18 by modeling ClyA as two series resistors (Eq. S19), ^96,116^ using the bulk NaCl conductivity at the given concentrations. Experimental errors (*n* = 3) were smaller than the symbol size and hence not shown. (b) Contour plot of the simulated (ePNP-NS) ionic conductance *G* = *I/V*_b_ as a function of *V*_b_ and *c*_s_. (c) Log-log plots of *G* as a function of *c*_s_ at +150 mV (top) and −150 mV (bottom)—comparing results obtained through experiments, PNP-NS and ePNP-NS simulations, and the simple resitor pore model. (d) Contour plot of the Na^+^ transport number 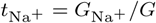, computed from the individual ionic conductances in the ePNP-NS simulation, as a function of *V*_b_ and *c*_s_. The 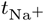 expresses the fraction of the ionic current is carried by Na^+^ ions, *i.e.*, the cation selectivity. (e) Simulated (PNP-NS and ePNP-NS) values of 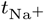 as a function of *c*_s_ for +150 mV (top) and −150 mV (bottom). Here, the ‘bulk’ line indicates the bulk NaCl cation transport number, represented by its empirical function 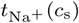 (see Tab. 1 and Tab. S2). The solid grey line represents 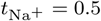.

The ability of a nanopore to conduct ions can be best expressed by its conductance: *G* = *I/V*_b_. We computed ClyA’s conductance with the ePNP-NS equations as a function of bias voltage (*V*_b_ = −200 to +200 mV) and bulk NaCl concentration (*c*_s_ = 0.005 to 5 M), of which a contour plot can be found in Fig. 2b. The near horizontal contour lines in the upper part of the plot (*c*_s_ > 1 M) show that, at high ionic strengths, ClyA maintains the same conductance regardless of the applied bias voltage. This behavior changes at intermediate concentrations (0.1 M < *c*_s_ < 1 M), where maintaining the same conductance level with increasing negative bias amplitudes requires increasing salt concentrations. Finally, at low salt concentrations (*c*_s_ < 0.1 M), the ionic conductance increases when reducing the negative voltage amplitude but subsequently levels out at positive bias voltages.

The cross-sections of the ionic conductance as a function of concentration at high positive and negative bias voltages (Fig. 2c), serve to demonstrate the differences between these respective regimes. The slope of the log-log plot at +150 mV is highly linear, suggesting that only a single mechanism of ionic conduction is at work. As touched upon above, both the bulk model and the ePNP-NS equations manage to capture the conductance at high ionic strengths, while only the latter performs well at low ionic strengths. The predictions made with the PNP-NS equations overestimate the conductance over the entire concentration range, but they do converge at very low ionic strenghts. At *V*_b_ = −150 mV, the log-log plot consists of two linear segments with a transition zone at *c*_s_ ≈ 0.15 M. This behavior is captured qualitatively by both simulation methods, but a perfect quantitative match is only found for the ePNP-NS equations. The bulk model exhibits the same single-sloped trend as seen at *V*_b_ = +150 mV.

The difference in ionic conduction at opposing bias voltages is also known as ionic current rectification (ICR): *α*(*V*_b_) = *G*(+*V*_b_)*/G*(−*V*_b_). ICR is a phenomenon often observed in nanopores that are both charged, and contain a degree of geometrical asymmetry along the central axis of the pore.^5,72,117^ As can be seen in Fig. S3, ClyA exhibits a strong degree of rectification, which is to be expected given its high negative charge (−72 *e* at pH 7.5) and different *cis* (≈3.3 nm) and *trans* (≈6 nm) entry diameters (Fig. 1a). We found *α* to increase monotoneously with the bias voltage magnitude, at least over the investigated range. We found the dependence of *α* on the ionic strength not to be monotonous, but rather rising rapidly to a peak value at ≈0.15 M, followed by a gradual decline towards unity at saturating salt concentrations.

The results and comparisons discussed above indicate that ClyA’s conductivity is dominated by the bulk electrolyte conductivity above physiological salt concentrations (*c*_s_ > 0.15 M). The breakdown of this simple dependency at lower ionic strengths is particularly evident at negative bias voltages and is likely caused by the overlapping of the electrical double layer inside the pore. This effectively excludes the co-ions—Cl^−^ in the case—from the interior of the pore, preventing them from contributing to total ionic conductance and resulting in a conductance dominated by the surface charge.^118^ The presence of only a single ion type inside the pore may also offer an explanation as to why the ePNP-NS equations falter at low ionic strengths. Because our ionic mobilities are derived from *bulk* ionic conductances, *i.e.*, for unconfined ions in a locally *electroneutral* environment, it is likely that our mobility model begins to break down under these conditions. ^119^ Another cause of the discrepancies could be a change in the shape or diameter of the nanopore at low salt concentrations, which cannot be captured by our simulation due to the static nature of its geometry and charge distributions. Nevertheless, our simplified 2D-axisymmetric model, in conjunction with the ePNP-NS equations, is able to accurately predict the ionic current flowing through ClyA for a wide range of experimentally relevant ionic strengths and bias voltages. This suggests that we managed to capture the essential physical phenomena that drive the ion and water transport through the nanopore both qualitatively and quantitatively. Hence, we expect the distribution of the resulting properties (*e.g.*, ion concentrations, ion fluxes, electric field and water velocity) to closely correspond to their real values.

#### Cation selectivity

The ion selectivity of a nanopore determines the preference with which it transports one ion type over the other. Experimentally, it is often determined by placing the pore in a salt gradient (*i.e.*, different salt concentrations in the *cis* and *trans* reservoirs) and measuring the potential at which the nanopore current is zero (reversal potential, *V*_r_).^96,101^ The Goldman-Hodkin-Katz (GHK) equation can then be used to convert *V*_r_ into the permeability ratio 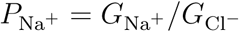. Here, we represent the ClyA’s ion selectivity (Figs. 2d and 2e) by the fraction of the total current that is carried by Na^+^ ions: the apparent Na^+^ transport number 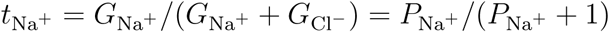.

Due its negatively charged interior, we found ClyA to be cation selective (*i.e.*, 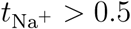) for all investigated voltages up to a bulk salt concentration of *c*_s_ ≈ 2 M NaCl (0.5 contour line in Fig. 2d). Above this concentraion, 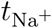 falls to a minimum of value of 0.45 at *c*_s_ ≈ 5 M, which is still ≈1.27 times its value in the bulk electrolyte (0.35). This indicates that—even at saturation—ClyA remains preferential towards cations. Below 2 M, the ion selectivity increases logarithmically with decreasing salt concentrations, but it also becomes more sensitive to the direction and magnitude of the electric field: with negative bias voltages yield higher ion selectivities (Fig. 2e). For example, to reach a selectivity of 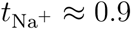, the salt concentration must fall to 0.05 M at +150 mV and to only 0.125 M at −150 mV.

Using the reversal potential method, Franceschini *et al.*^101^ found ClyA’s ion selectivity to be 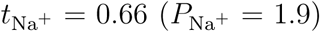. In our ePNP-NS simultation, this corresponds the selectivity at *c*_s_ = 0.5 M(at *V*_b_ = 0 mV), a value that lies in between the *cis* 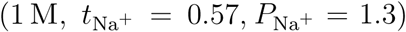 and *trans* 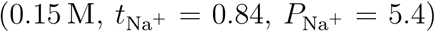) reservoir concentrations used in their experiment. Even though measuring the reversal potential can give valuable insights into the selectivity ion channels and small nanopores, it should be used with caution for larger nanopores such as ClyA. The GHK equation does not consider the ionic flux due to the electro-osmotic flow and assumes that the Nernst-Einstein relation holds for all used concentrations. These two effects should not be ignored as they contribute significantly to the nanopore’s total conductance. Furthermore, because the ion selectivity depends strongly on the ionic strength, the measured reversal potential will necessarily be influenced by the chosen salt gradient and represent the selectivity at an undetermined intermediate concentration.

### Ion Concentration Distribution

Following the validation of the model in preceding section, we now proceed by describing the local ionic concentrations inside ClyA. Detailed knowlegde of the ionic environment can be valuable to experimentalists who seek to trap and study single enzymes with ClyA. ^30,97,100^ Moreover, it gives insight into the origin of the ion current rectification, ion selectivity and the electro-osmotic flow. We evaluated the densities of both Na^+^ and Cl^−^ ions inside ClyA and investigated the effect of bulk salt concentration and the bias voltage on 1) the distribution of their concentration (Figs. 3a to 3c) and 2) the accumulation of mobile charges inside the pore as a result of their asymmetric enhancement or depletion (Figs. 3d to 3f).

**Figure 3.**
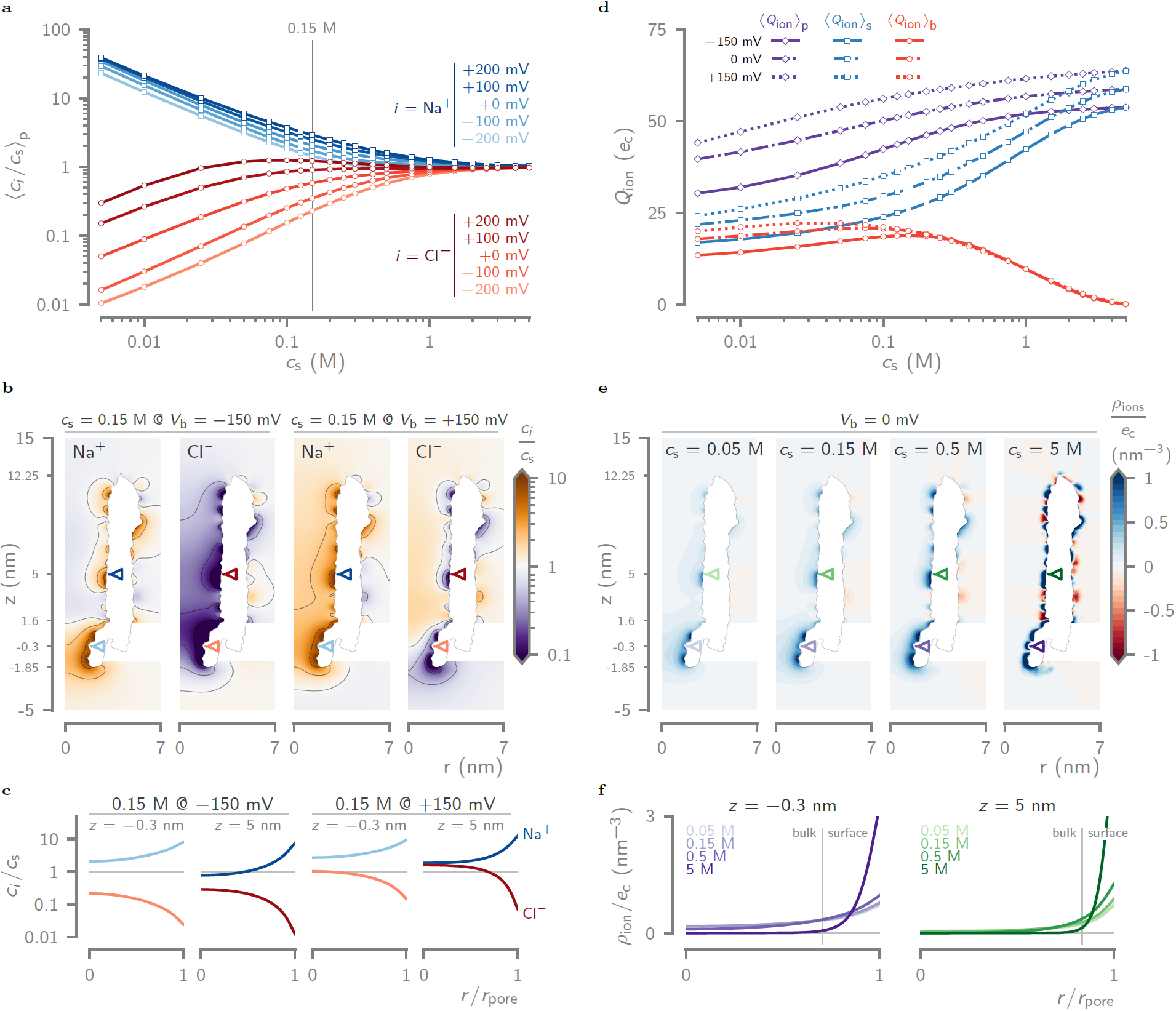
Ion concentration distribution inside ClyA. (a) Relative Na^+^ and Cl^−^ concentrations averaged over the entire pore volume (⟨*c*_*i*_*/c*_s_⟩_p_) as a function of the reservoir salt concentration (*c*_s_ = 0.005 to 5 M) and bias voltage (*V*_b_ = −200 to +200 mV). (b) Contour plots of the relative ion concentration (*c*_*i*_*/c*_s_) for both Na^+^ and Cl^−^ for *c*_s_ = 0.15 M and at *V*_b_ = −150 and +150 mV. (c) The relative Na^+^ and Cl^−^ concentration profiles along the radius of the pore, through the middle of the constriction (*z* = −0.3 nm) and the *lumen* (*z* = 5 nm), as indicated by the arrows in (b). (d) The average number of ionic charges inside the pore ⟨*Q*_ion_⟩_p_, is distributed between the those close to the pore’s surface ⟨*Q*_ion_⟩_s_, *i.e.*, within 0.5 nm of the wall, and those in the ‘bulk’ of the pore’s interior ⟨*Q*_ion_⟩_b_. (e) Cross-section contour plots of the ion space charge density (*ρ*_ion_), expressed as number of elementary charges per nm^3^, at *V*_b_ = 0 mV and for *c*_s_ = 0.05, 0.15, 0.5 and 5 M. (f) Radial cross-sections of the *ρ*_ion_ at the center of the constriction (*z* = −0.3 nm) and the *lumen* (*z* = 5 nm) of ClyA. The vertical line represents the the division between ions in the ‘bulk’ (*d* > 0.5 nm) of the pore and those located near its surface (*d* ≤ 0.5 nm).

#### Relative cation and anion concentrations

The relative ion concentration averaged over the entire volume of the pore (⟨*c*_*i*_*/c*_s_⟩_p_) as a function of bulk concentration (Fig. 3a), gives a good measure for global situation inside the pore. At low ionic strengths (*c*_s_ < 0.05 M), our simulation predicts a strong enhancement of the Na^+^ (cation) concentration 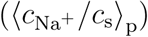 and a clear depletion of the Cl^−^ (anion) concentration 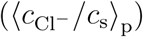 inside the pore. This effect, however, dimishes rapidly with increasing ionic strengths, which can be explained by the electrolytic screening of the negative charges lining the walls of ClyA (*i.e.*, the electrical double layer). At low bulk concentrations the number of ions in the bulk is sparse, leading to the attraction and repulsion of respectively as many Na^+^ and Cl^−^ ions as the chemical potential allows. As the concentration increases, the overall availibility of ions improves and the extreme concentration differences between the pore and the bulk are no longer required to offset ClyA’s fixed charges. For example, changing the bulk concentration from 0.005 to 0.05 M under equilibrium conditions (*V*_b_ = 0 mV), causes the relative Na^+^ and Cl^−^ concentrations to fall and rise from 34 to 4.4 and from 0.05 to 0.31, respectively. At *c*_s_ = 0.15 M, the values still differ significantly from bulk(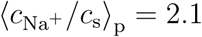 and 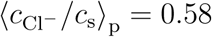), but they fall back to <15 % of unity at *c*_s_ = 1 M (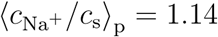 and 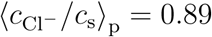).

We also observed a stark difference between their sensitivities to the applied bias voltage, particularly at low salt concentrations (Fig. 3a, left of <0.15 M line). Whereas the Na^+^ concentration shows only a limited response, the Cl^−^ concentration changes much more dramatically. For example, at *c*_s_ = 0.15 M and from *V*_b_ = −150 to +150 mV, 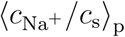 rises ≈1.7-fold (1.6 to 2.7) and 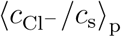 increases ≈3.8-fold (0.28 to 1.06). This difference is clearly visualized by the contour plots of the relative ion concentrations (*c*_*i*_*/c*_s_) at *c*_s_ = 0.15 M and for *V*_b_ = *text*−150*and*+150 mV (Fig. 3b). They reveal that the *trans* constriction (−1.85 < *z* < 1.6 nm) remains depleted of Cl^−^ and enhanced in Na^+^ for both *V*_b_ = −150 mV and *V*_b_ = +150 mV. This is not the case in the *lumen* (1.6 < *z* < 12.25 nm), in which the Na^+^ concentration is bulk-like for *V*_b_ < 0 mV and enhanced for *V*_b_ > 0 mV. Conversely, the number of Cl^−^ ions becomes more and more depleted in the *lumen* for increasing negative bias magnitudes, and it is virtually bulk-like at higher positive bias voltages. This is further exemplified by the radial profiles of the ion concentrations (Fig. 3c) through the middle of the constriction (*z* = −0.3 nm) and the *lumen* (*z* = 5 nm), which also clearly shows the extent of the electrical double layer.

#### Ion Charge Density

The formation of an electrical double layer inside the pore, and the resulting asymmetry in the cation and anion concentrations, gives rise to a net charge density inside the pore (*ρ*_ion_, Eq. 2). To investigate the distribution of these charges within ClyA, we divided the interior volume of the pore into a ‘surface’ and a ‘bulk’ region. The bulk region encompasses a cylindrical volume at the center of the pore up until 0.5 nm distance from the wall (*d* ≥ 0.5 nm), and the surface region includes the remaining volume between the bulk and the nanopore wall (*d* < 0.5 nm). Integration of *ρ*_ion_ over the ‘surface’ and ‘bulk’ regions yields the average number of mobile charges present inside those locations (Fig. 3d): ⟨*Q*_ion_⟩_b_ and ⟨*Q*_ion_⟩_s_, respectively. Although the total number of charges inside the pore, ⟨*Q*_ion_⟩_p_ = ⟨*Q*_ion_⟩_b_ + ⟨*Q*_ion_⟩_s_, rises appreciatively with increasing reservoir concentrations, the majority of these additional charges are confined to the surface of the pore. Up until *c*_s_ = ≈0.15 M, ⟨*Q*_ion_⟩_p_ is distributed equally between the surface (≈ + 27 *e*) and bulk (≈ + 22 *e*) layers. At higher concentrations, the number of charges in the surface layer more than doubles (towards ⟨*Q*_ion_⟩_s_ = +58 *e* at 5 M), and those in the bulk of the pore diminish (towards ⟨*Q*_ion_⟩_b_ ≈ 0 *e* at 5 M). The total number of mobile charges ⟨*Q*_ion_⟩_p_ also depend on the applied bias voltage. As can be seen from our simulation results for three different voltages (Fig. 3d), ⟨*Q*_ion_⟩_p_ is approximately +10 to +15 *e* higher at *V*_b_ = +150 mV as compared to *V*_b_ = −150 mV for the full range of ion concentrations. Interestingly, at higher salt concentrations this increased number of charges with voltage is fully accounted for by the surface charges, as the charge density in the bulk does not change with the applied voltage.

The cross-section contour plots of *ρ*_ion_ inside ClyA for four different bulk concentrations (*c*_s_ = 0.05, 0.15, 0.5 and 5 M) reveal the redistribution of the mobile charges with increasing ionic strength in more detail. Up until a bulk concentration of *c*_s_ = ≤0.5 M, the electrical double layer inside the pore overlaps significantly with itself (Fig. 3e, 3 leftmost panels), giving rise to a net positive charge spread diffusively throughout the entire pore. Moreover, the absence of Cl^−^ ions prevents the formation of any significant negative charge densities next to the few positively charged residues lining the pore walls. The situation at high salt concentrations (*e.g.*, 5 M) is very different, with almost no charge density within the ‘bulk’ of the pore *lumen* (*d* ≥ 0.5 nm), but with pockets of highly charged and alternating positive and negative charge densities close to the nanopore wall (Fig. 3e, rightmost panel). This sharp confinement is shown clearly by the radial density profiles (Fig. 3f) drawn through the constriction (*z* = −0.3 nm, purple triangles) and the *lumen* (*z* = 5 nm, green triangles).

### Electrostatic Potential and Energy

The electrostatic potential, or rather the change thereof, is one of the primary driving forces within a nanopore. Typically, the potential can be split into an intrinsic, ‘equilibrium’ component and an external, ‘non-equilibrium’ contribution. The latter is also known as the bias voltage applied between the *trans* and the *cis* reservoirs, and most of it changes across the short distance of the nanopore itself. For a 14 nm-long nanopore such as ClyA, a potential drop of 10 to 100 mV can easily create an electric field of 10^6^–10^7^ V · m^−1^ inside the pore. The intrinsic electrostatic potential results from the charged residues embedded in the interior walls of the protein and can significantly influence the transport of ions and water molecules through the pore.^46,48,57,120^ In the following section we aim to describe the most salient features of this potential and the extent of its influence over the entire investigated concentration range (Fig. 4). Moreover, because electromigration is the primary contributor to the ionic current, a proper understanding of the electrostatic landscape—and the energy barriers faced by the ions traversing the pore—should provide a more quantitative explanation for the origin of ClyA’s rectification, ion selectivity and asymmetric ion concentrations.

**Figure 4.**
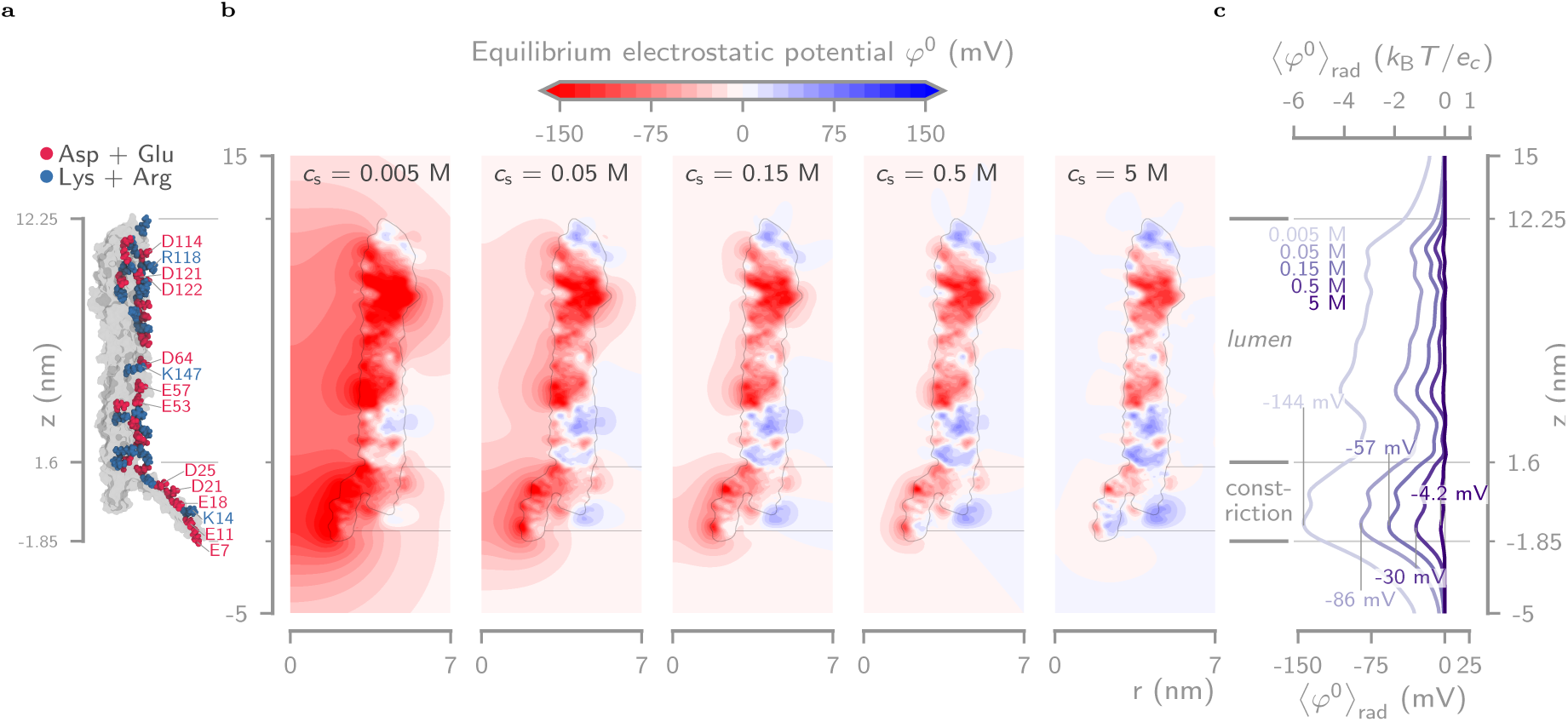
Equilibrium electrostatic potential inside ClyA. (a) A single subunit of ClyA in which all amino acids with a net charge and whose side chains face the inside of the pore, *i.e.*, those contribute the most to the electrostatic potential, are highlighted. Negatively (Asp+Glu) and positively and positively (Lys+Arg) charged residues are colored in red and blue, respectively. (b) As a result of these fixed charges ClyA exhibits a complex electrostatic potential (*ϕ*) landscape at equilibrium (*i.e.*, at *V*_b_ = 0 mV), and whose values inside the pore we have plotted for several key concentrations (*c*_s_ = 0.005, 0.05, 0.15, 0.5 and 5 M). Note that even at physiological salt concentrations (*c*_s_ = 0.15 M), the negative electrostatic potential extends significantly inside the *lumen* (1.6 < *z* < 12.25 nm), and even more so inside the *trans* constriction (1.85 < *z* < 1.6 nm). For the former, localized influential negative ‘hotspots’ can be found in the middle (4 < *z* < 6 nm) and at the *cis* entry (10 < *z* < 12 nm). (c) Radial average of the equilibrium electrostatic potential along the length of the pore (⟨*ϕ*⟩_rad_) for the same concentrations as in (b). Even though the *lumen* of the pore is almost fully screened for *c*_s_ > 0.5 M, the constriction still retains some of its negative influence even at 5 M.

#### A few important charged residues

The interior walls of the ClyA nanopore (Fig. 4a) are riddled with negatively charged amino amids (*i.e.*, aspargine or glutamine), interspaced by a few positively charged residues (*i.e.*, lysine or arginine). When grouping these charges by proximity, we found three clusters with significantly more negative than positive residues: inside the *trans* constriction (−1.85 < *z* < 1.6 nm; E7, E11, K14, E18, D21, D25), in the middle of the *cis lumen* (4 < *z* < 6 nm; E53, E57, D64, K147) and at the top of the pore (10 < *z* < 12 nm; D114, R118, D121, D122). As we shall see, these clusters leave strong negative fingerprints in the global electrostatic potential.

#### Distribution of the equilibrium electrostatic potential

The electrostatic potential at equilibrium (*φ*^0^, *i.e.*, at *V*_b_ = 0 mV) reveals the effect of ClyA’s fixed charges on the potential inside the pore (Fig. 4b). Due to electric screening by the mobile charge carriers in the electrolyte, however, the extent of their influence strongly depends on the bulk ionic strength. The contour plot cross-sections of *φ*^0^ for *c*_s_ = 0.005, 0.05, 0.15, 0.5 and 5 M (Fig. 4b) and their corresponding radial averages (Fig. 4c) demonstrate this effect aptly. The radial average (⟨*φ*^0^ ⟩_rad_) represents the mean value along the longitudunal axis of the pore and can be computed using

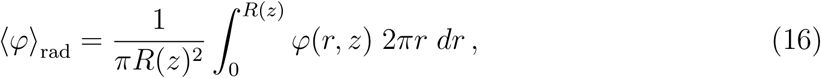

where *R*(*z*) is taken as the pore radius for −1.85 ≤ *z* ≤ 12.25 nm, 2 nm for *z* < −1.85 nm and 4 nm for *z* > 12.25 nm. Starting from the *cis* entry (*z* ≈ 10 nm), the electrostatic potential is dominated by the acidic residues D114, D121 and D122, resulting in a rapid reduction of ⟨*φ*^0^⟩_rad_ upon entering the pore. Next, ⟨*φ*^0^⟩_rad_ slowly descreases up until the middle of the *lumen* (*z* ≈ 5 nm), where the next set of negative residues, namely E53, E57 and D64, lower it even further. After a brief increase, ⟨*φ*^0^⟩_rad_ attains its maximum amplitude inside the *trans* constriction (*z* ≈ 0 nm) due to the close proximity of the amino acids E7, E11, E18, D21 and D25, and then quickly falls to 0 inside the *trans* reservoir.

At low ionic strengths (*c*_s_ < 0.05 M), the lack of sufficient ionic screening results in relatively high negative potentials throughout the entire pore. For example, for *c*_s_ = 0.005 and 0.05 M, the ⟨*φ*^0^⟩_rad_ inside the constriction ramps up to values of −144 mV (−5.60 k_B_T*/*e) and −86 mV (−3.35 k_B_T*/*e), respectively. These values significantly exceed the single ion thermal voltage *k*_B_*T/e* = 25.7 mV. Hence, on the one hand they prohibit anions such as Cl^−^ from entering the pore, and on the other they attract cations such as Na^+^ and trap them inside the pore. For intermediate concentrations (0.05 ≤ *c*_s_ < 0.5 M) the influence of the negative charges becomes increasingly confined to several ‘hotspots’ near the nanopore walls, most notably at entry of the pore (10 < *z* < 12 nm), in the middle of the *lumen* (4 < *z* < 6 nm), and in the constriction (−1.85 < *z* < 1.6 nm), in accordance with the charge groups discussed in the previous section. Even though the magnitude of ⟨*φ*^0^⟩_rad_ at *c*_s_ = 0.15 M drops below 1 k_B_T*/*e inside the *lumen* (⟨*φ*^0^⟩_rad_ ≈ −14 mV), it remains strongly negative inside the constriction (⟨*φ*^0^⟩_rad_ ≈ −47 mV). Finally, at high concentrations (*c*_s_ ≥ 0.5 M) the potential is close to ≈0 mV over the entire *lumen* of the pore, with only a small negative potential remaining inside the constriction. A summary of the most salient ⟨*φ*⟩)_rad_ values can be found in Tab. S3.

#### Non-equilibrium electrostatic energy at +150 and −150 mV

To link back the observed ionic conductance properties to the electrostatic potential, we computed the radially averaged electrostatic energy for a monovalent ion, ⟨*U*_E,*i*_⟩_rad_ = *z*_*i*_*e* ⟨*φ*⟩_rad_, at *V*_b_ = +150 −150mV for the entire range of simulated ionic strengths (Fig. 5a). The resulting energy plot represents the energy landscape—filled with barriers (hills) or traps (valleys)—that a positive or negative ion must traverse in order to contribute positively to (*i.e.*, increase) the ionic current.

**Figure 5.**
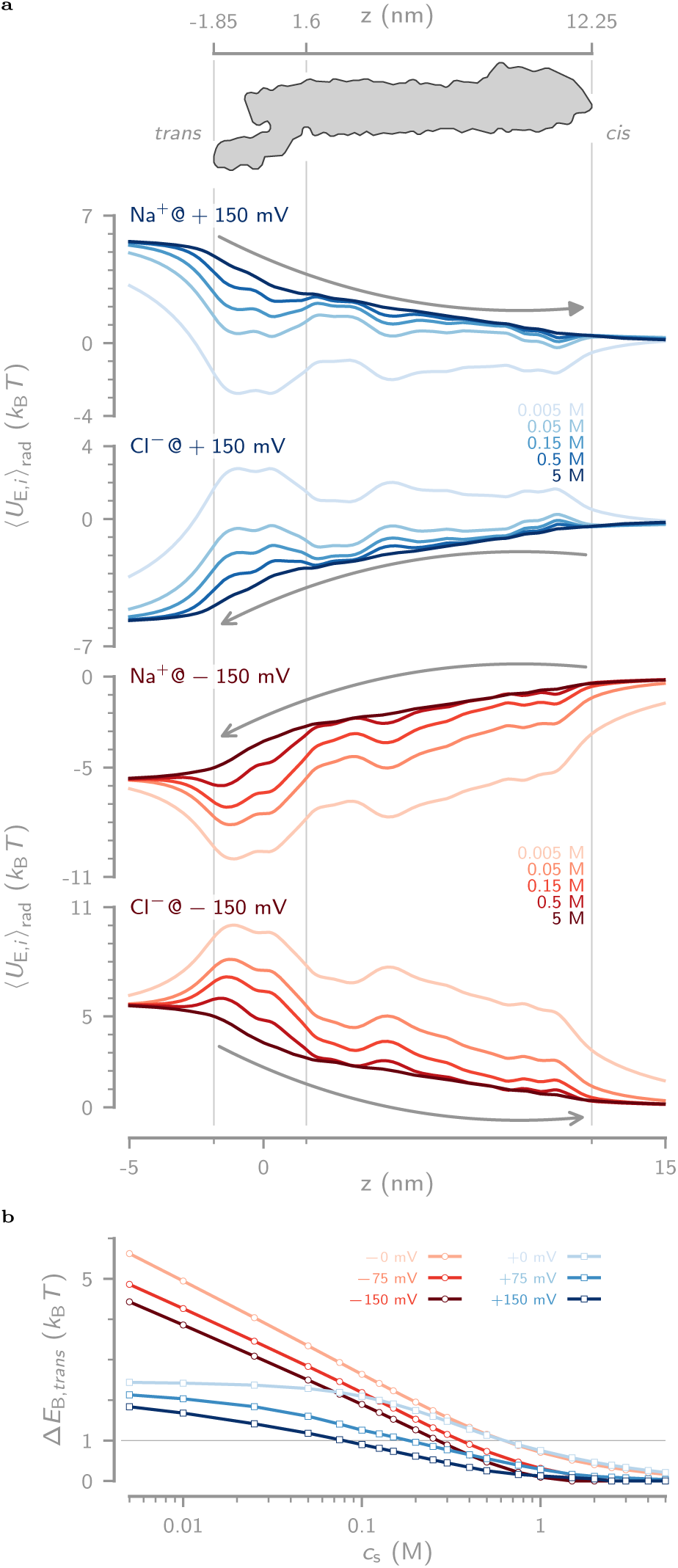
Non-equilibrium electrostatic energy landscape for single ions. (a) Radially averaged non-equilibrium electrostatic energy landscape for single ions, ⟨*U*_E,*i*_⟩_rad_ = *z*_*i*_*e* ⟨*ϕ*⟩_rad_, as calculated directly from the radial electrostatic potential at *V*_b_ = +150 and −150 mV for monovalent cations and anions. The grey arrows indicate the direction in which the ions must travel in order for them to contribute positively to the ionic current. (b) Height of the electrostatic energy barrier (Δ*E*_B,*i*_) at the *trans* constriction as a function of the bulk salt concentration. Note that Δ*E*_B,*i*_ is much higher for negative voltages and rises logarithmically at lower concentrations. The divergence between +0 mV and −0 mV for *c*_s_ < 0.3 M highlights the difference in barrier height when traversing the pore from *cis* to *trans* or *vice versa*.

At positive bias voltages, cations traverse the pore from *trans* to *cis* (Fig. 5a, first plot). Upon entering the negatively charged constriction, their electrostatic energy drops dramatically, followed by a relatively flat section with a small barrier for entry in the *lumen* at *z* ≈ 1.6 nm. At very low ionic strengths (*c*_s_ < 0.05 M), the energy at *trans* is significantly lower than the energy of the cation in the *cis* compartment (*e.g.*, Δ ⟨*U*_E,*i*_⟩_rad_ > 2 k_B_T at 0.005 M), forcing the ions to accumulate inside the pore. At higher concentrations (*c*_s_ > 0.05 M), the increased screening smooths out the potential drop inside the pore, allowing the cations to migrate unhindered across the entire length of the pore. Anions at *V*_b_ = +150 mV travel from *cis* to *trans* (Fig. 5a, second plot) and must overcome energy barriers at both sides of the pore. The former prevents anions such as Cl^−^ from entering the pore. However, because its magnitude is attenuated strongly with increasing salt concentration (from ≈1.7 k_B_T at *c*_s_ = 0.005 M to ≈0.5 k_B_T at *c*_s_ = 0.05 M), it is only relevant at lower ionic strengths (*c*_s_ < 0.05 M). Once inside the *lumen*, anions can move relatively unencumbered to the *trans* constriction, where they face the second, more significant energy barrier. This prevents them from fully translocating and causes them to accumulate inside the lumen, explaining the higher Cl^−^ concentrations at positive bias voltages. As with the cations, an increase in the ionic strength significantly reduces these hurdles, resulting in a much smoother landscape for *c*_s_ > 0.15 M.

At negative voltages, cations move through the pore from *cis* to *trans*, with a slow and continuous drop of the electrostatic energy throughout the *lumen* of the pore up until the constriction (Fig. 5a, third plot). This results in the efficient removal of cations from the pore *lumen*, and explains the lower Na^+^ concentration observed at positive voltages (Fig. 3a). To fully exit from the pore, however, cations must overcome a large energy barrier, which reduces the nanopore’s ability to conduct cations compared to positive potentials and hence contributes to the ion current rectification. The situation for anions at negative bias voltages (*i.e.*, travelling from *trans* to *cis*) is very different (Fig. 5a, fourth plot). Their ability to even enter the pore is severely hampered by an energy barrier of a few k_B_T at the *trans* constriction. Any anions that do cross this barrier, and those still present in the *lumen* of ClyA, will be rapidly move towards the *cis* entry and exit from the pore due to a continuous drop of electrostatic energy. This effectively depletes the entire *lumen* of anions, which can be observed from the much lower Cl^−^ concentrations at negative voltages (see Fig. 3a).

#### Concentration and voltage dependencies of the energy barrier at the constriction

Many biological nanopores contain constrictions that play crucial roles in shaping their ionic conductance properties. ^14,28,101^ The reason for this is two-fold, 1) the narrowest part dominates the overall resistance of the pore and 2) confinement of charged residues results in much larger electrostatic energy barriers. With its highly negatively charged *trans* constriction, ClyA’s affinity for transport of anions is diminished and that for cations is enhanced compared to bulk, even at high ionic strengths (Fig. 2e). ^96^ To further elucidate the significance of the *trans* electrostatic barrier (Δ*E*_B,*i*_), we quantified its height at positive and negative voltages as a function of the salt concentration (Fig. 5b).

Due to the tilting of the nergy landscape, the application of a bias voltage lowers the magnitude of the energy barrier for all conditions. Likewike, increasing the bulk salt concentration results in a continuous decrease of Δ*E*_B,*i*_. For *c*_s_ > 0.5 M, Δ*E*_B,*i*_ falls below 1 k_B_T regardless of the bias voltage, reducing its effect on the ion transport through the pore. The barrier heights for ions under the influence of a positive bias voltage (*i.e.*, Na^+^ moving from *trans* to *cis* and Cl^−^ moving from *cis* to *trans*), experience a Δ*E*_B,*i*_ roughly half that of those under a negative voltage. For example, upon increasing the salt concentration from 0.005 to 0.15 M, Δ*E*_B,*i*_ dropped from 1.8 to 0.73 k_B_T at *V*_b_ = +150 mV and from 4.4 to 1.5 k_B_T at *V*_b_ = −150 mV. This explains the differences in ion selectivity observed between positive and negative bias voltages (Fig. 2e). At *c*_s_ = 0.15 M, for example, the ion selectivity 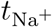 increases from 0.79 at *V*_b_ = +150 mV to 0.88 at *V*_b_ = −150 mV.

### Transport of Water Through ClyA

The charged nature of many nanopores gives rise to a net flux of water through the pore, called the electro-osmotic flow (EOF).^37,82,121^ The EOF not only contributes significantly to the ionic current, but the magnitude of the viscous drag force it exerts is often of the same order as the Coulombic electrophoretic force (EPF). Hence, it strongly influences the capture and translocation of biomolecules including nucleic acids,^36,47,122^ peptides, ^28,123,124^ and proteins.^25,27,30,96–99,125^ Because the drag exerted by the EOF depends primarily on the size and shape of the biomolecule of interest and not on its charge,^125^ it can be harnassed to capture molecules even against the electric field. ^25^ The EOF is a consequence of interaction between the fixed charges on the nanopore walls and mobile charges in the electrolyte. It can be described by two closely related mechanisms: 1) the excess transport of the hydration shell water molecules in one direction due to the pore’s ion selectivity, and 2) the viscous drag exerted by the unidirectional movement of the electrical double layer inside the pore. The first mechanism likely dominates in pores with a diameter close to that of the hydrated ions (≤1 nm) such as αHL or FraC,^28,124^ while the second is expected to be stronger for larger pores (>1 nm), such as ClyA^25,125^ or most solid-state nanopores.^37,39^ In our simulation, the EOF follows the coupling of the Navier-Stokes and the Poisson-Nernst-Planck equations through a volume force term (Eq. 13), which dictates that the electric field exerts a net force on the fluid if it contains a net charge density—as is the case for the electrical double layer lining the walls of ClyA (Fig. 3d). In the following section we aim to quantitatively and a qualitatively describe the properties of the water velocity inside ClyA and how it is influenced by the bulk ionic strength and the applied bias voltage (Fig. 6).

**Figure 6.**
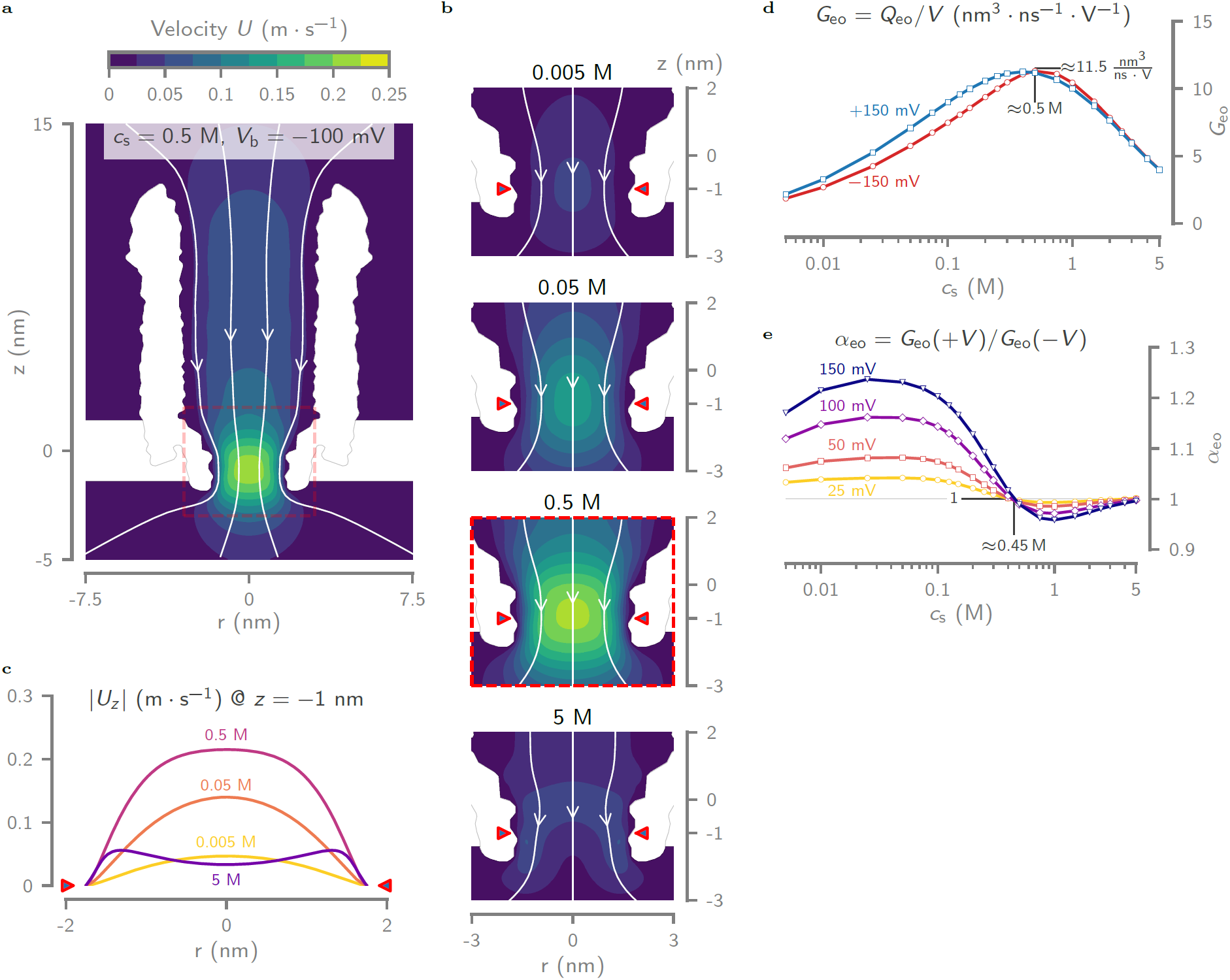
Concentration and voltage dependency of the electro-osmotic flow inside ClyA. (a) Contour plot of the electro-osmotic flow (EOF) velocity ***u*** at 0.5 M and −100 mV bias voltage. The arrows on the streamlines indicate the direction of the flow. As observed experimentally ^96^ and expected from a negatively charged conical nanopore, the EOF follows the direction of the cation, *i.e.*, from *cis* to *trans* under negative bias voltages and *vice versa* for positive ones. (b) Contour plots of the EOF field in the *trans* constriction for various salt concentrations at *V*_b_ = −100 mV and (c) cross-sections of the absolute value of the water velocity |*U*_*z*_| at *z* = −1 nm. Notice that at high salt concentrations (*c*_s_ > 1 M), the velocity profile exhibits two ‘lobes’ close to the nanopore walls and hence deviates from the parabolic shape observed at lower ionic strengths. (d) Concentration dependence of the electro-osmotic conductance *G*_eo_ = *Q*_eo_*/V*_b_, with *Q*_eo_ the total flow rate through the pore (Eq. 21). In the low concentration regime, *G*_eo_ increases rapidly between 0.005 and 0.5 M after which it decreases logarithmically for higher concentrations. (e) The rectification of the electro-osmotic conductance (*α*_eo_(−*V*) = *G*_eo_(+*V*)*/G*_eo_(−*V*)) plotted against the bulk salt concentration. The *α*_eo_ increases with bias voltage and exhibits an inversion point at *c*_s_ *≈* 0.45 M.

#### Direction, magnitude and distribution of the water velocity

As expected, given ClyA’s negatively charged interior surface and the resulting positively charged electrical double layer, the direction of the net water flow inside ClyA follows the electric field, *i.e.*, from the *cis* to the *trans* at negative bias voltages (Fig. 6a). This corresponds to the observations and analysis of single-molecule protein capture^96^ and trapping^97,98,125^ experiments using the ClyA-AS nanopore. Along the longitudinal axis (*z*) of the pore, the water velocity is governed by the conservation of mass, meaning it is lowest in the wide *cis lumen* and highest in the narrow *trans* constriction (Fig. 6a). For example, at *V*_b_ = −100 mV and *c*_s_ = 0.5 M the velocity at the center of the pore is ≈0.07 m · s^−1^ in the *lumen* and ≈0.21 m · s^−1^ in the constriction.

Along the radial axis (*r*), ***u*** has a parabolic profile with the highest value at the center of the pore and the lowest at the wall due to the no-slip boundary condition (Figs. 6b and 6c). Such a parabolic profile contrasts the expected ‘plug flow’ for an EOF, but follows logically from the overlap of the electrical double layer inside the pore and the resulting uniform volume force—analogous to a gravity- or pressure-driven Stokes flow. At concentrations higher than 0.5 M, however, the increasing degree of confinement of the double layer—and its charge—to the nanopore walls (see Fig. 3d, ⟨*Q*_ion_⟩_s_) results in a flattening of the central maximum and hence a plug flow profile. Interestingly, at very high salt concentrations (*c*_s_ ≥ 1 M) the velocity profile in the constriction exhibits a dimple at the center of the pore. This is the result of a self-induced pressure gradient caused by the expansion of the EOF as it exits the pore.^126^

#### Influence of bulk ionic strength and bias voltage on the electro-osmotic conductance

In analogy to the ionic conductance, the amount of water transported by ClyA can be expressed by the electro-osmotic conductance *G*_eo_ = *Q*_eo_*/V*_b_ (Fig. 6d). Here, *Q*_eo_ is the net volumetric flow rate of water through the pore and computed by integrating the water velocity across the reservoir boundary (Eq. 21). The strength of the EOF depends strongly and non-monotonically on the bulk ionic strength: *G*_eo_ rapidly increases with ionic strength until a peak value is reached at *c*_s_ ≈ 0.5 M, followed by a gradual logarithmic decline (Fig. 6d). For example, at *V*_b_ = −150 mV, *G*_eo_ changes from 1.85, 11.3 to 4.00 nm^3^ · ns^−1^ · V^−1^ at for *c*_s_ = 0.005, 0.5 and 5 M, respectively.

The sensitivity of the EOF to the magnitude and sign of the bias voltage is given by the electro-osmotic conductance rectification *α*_eo_(*V*) = *G*_eo_(+*V*)*/G*_eo_(−*V*) (Fig. 6e). For all voltages magnitudes, *α*_eo_ shows a maximum at *c*_s_ ≈ 0.045 M, after which it falls rapidly to reach unity (*α*_eo_ = 1) at approximately *c*_s_ ≈ 0.45 M. A minimum is then reached at ≈1 M, followed by a gradual approach towards unity at *c*_s_ = 5 M.

## Conclusions

We have developed an extended version of the Poison-Nernst-Planck-Navier-Stokes (ePNP-NS) equations and used it to model the transport of ions and water through the biological nanopore ClyA. Our ePNP-NS equations combine many of the improvements to the PNP-NS equations available in literature, in addition to several new corrections. These include the finite size of the ions, self-consistent concentration- and positional-dependent parametrization of the ionic transport coefficients (diffusion coefficient and mobility) and of the electrolyte properties (density, viscosity and relative permittivity).

We have verified our approach by matching experimental results to a very high degree of accuracy and the model parameters where gauged on other experiments, ultimately leaving no degrees of freedom. This shows that a continuum approach to modeling biological nanopores is not only feasible but to a very high degree predictive. We are confident that such a model as presented in this paper, constitutes a highly practical tool that can help with 1) elucidating the link between ionic current observed during a nanopore experiment and the actual physical phenomenon, 2) describing the electrophoretic and electro-osmotic properties of any biological nanopore and 3) guiding the design of new variants of existing nanopores.

In the analysis of the operation of ClyA with our model we found that the ionic currents depend strongly on the details of the electric field structure within the pore which define potential barriers for the ion species. These barriers in turn are influenced by both the bulk ionic strength and the applied bias voltage. Nevertheless, despite its simplified 2D-axisymmetric nature, our model is able to obtain meaningful and predictive results in terms of the ionic current, the ion concentrations, the electrostatic potential and electro-osmotic flow velocity. Some inaccuracies still arise at low salt concentrations, but these can likely be resolved with an improved mobility model.

Although not explicitly investigated in the current work, there are no obvious reasons why the model framework is not directly applicable to other biological nanopores, solid-state nanopores and other nanoscale ionic transport problems. These systems can be treated analogously and their properties could be mapped out systematically using our ePNP-NS model, greatly enhancing the understanding and interpretation of experimental results.

## Materials and Methods

### ClyA-AS homology model

A full atom model of ClyA-AS^96^ was built and optimized (MODELLER v9.18^127^) by introduction of the following point mutations in each of the 12 chains of the wild-type ClyA crystal structure (PDBID: 2WCD^95^): K8Q, N15S, Q38K, A57G, T67V, C87A, A90V, A95S, L99Q, E103G, K118R, L119I, I124V, T125K, V136T, F166Y, K172R, V185I, K212N, K214R, S217T, T224S, N227A, T244A, E276G, C285S, K290Q. Next, the conformation of all mutated side chains was optimized with an double annealing protocol (heating: 150, 250, 400, 700 and 1000 K, cooling: 1000, 800, 600, 500, 400 and 300 K) where at each temperature the energy was minimized for 200 iterations with a conjugate gradients algorithm (4 fs timestep).^128^ The first anneal was performed solely on the mutated residues themselves, and the second run also took the non-bonded interactions with the neighboring atoms into account. The refined nanopore structure was then embedded in the center of an 18 nm×18 nm equilibrated DPhPC lipid bilayer patch by manual removal of all overlapping lipids, resulting in 463 lipid molecules. The bilayer was created with the CHARMM-GUI^129^ membrane builder^130^ and equilibrated with NAMD^131^, as described in detail in ref. [132]. The system was then solvated in a box of 18 nm×18 nm×32 nm by addition of 214640 TIP3 water molecules (VMD solvate plugin), and the global charge was neutralized by replacing 1276 random water molecules with 674 Na^+^ and 602 Cl^−^ ions (VMD autoionize plugin).^102^

#### Molecular dynamics simulations

Using molecular dynamics (MD) with NAMD 2.12 (2 fs timestep, CHARMM36 forcefield^133^), the final system was minimized for 5 ps, heated from 0 to 298.15 K in 4 ps and equilibrated for 4 ns as NpT ensemble. ^46^ Finally a 30 ns production run was performed using a NVT ensemble at 298.15 K and the atomic coordinates saved every 5 ps. Note that structural deterioration was prevented by harmonically restraining the protein’s C_*α*_ atoms to their original positions (spring constant of 695 pN · nm^−1^) during all MD runs.^48^

#### Axially symmetric geometry

The 2D-axisymmetric geometry of the ClyA-AS nanopore (Fig. 1c) was derived directly from its full atom model by radially averaging the molecular density. Briefly, 50 sets of atomic coordinates were extracted from the final 5 ns of the coordinates of the 30 ns MD production run (*i.e.*, every 100 fs) and aligned by minimizing the RMSD between their backbone atoms (VMD RMSD tool). Next, we computed and averaged the 3D-dimensional molecular density maps of all 50 structures on a 0.5 Å resolution grid using the Gaussian function^104^

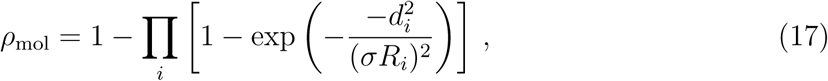

where for each atom *i, R*_*i*_ is its Van der Waals radius, 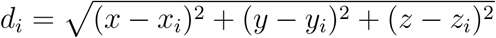 is the distance of grid coordinates (*x, y, z*) from the atom center (*x*_*i*_, *y*_*i*_, *z*_*i*_) and *σ* = 0.93 is a width factor. The resulting 3D density map was then radially averaged along the z-axis, relative to the center of the pore to obtain a 2D-axisymmetric density map. The contourline at 25 % density was used as the nanopore simulation geometry, after manual removal of overlapping and superfluous vertices to improve the quality of the final computational mesh.

#### Axially symmetric charge density

The 2D-axially symmetric charge distribution (Fig. 1d was also derived directly from the 50 sets of aligned nanopore coordinates) that were used for the geometry. Inspired by how charges are represented in the particle mesh Ewald (PME) method,^46^ we computed the fixed charge distribution of the nanopore 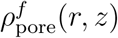 by assuming that an atom *i* of partial charge *δ*_*i*_, at the location (*x*_*i*_, *y*_*i*_, *z*_*i*_) in the full 3D atomistic pore model, contributes an amount *δ*_*i*_*/*2*πr*_*i*_ to the partial charge at a point (*r*_*i*_, *z*_*i*_) with 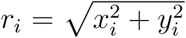 in the averaged 2D-axisymmetric model. This effectively spreads the charge over all angles to achieve axial symmetry. We assumed a Gaussian distribution of the space charge density of each atom *i* around its respective 2D-axisymmetric coordinates (*r*_*i*_, *z*_*i*_) is

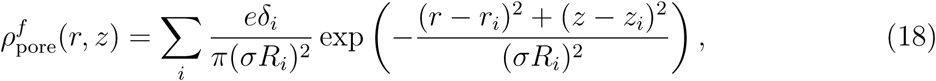

where *R*_*i*_ is the atom radius, *σ* = 0.5 is the sharpness factor and *e* is the elementary charge. To embed 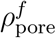 with sufficient detail, yet efficiently, into a numeric solver, the spatial co-ordinates were discretized with a grid spacing of 0.005 nm in the domain of 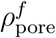 and pre-computed values were used during the solver runtime. All partial charges (at pH 7.5) and radii were taken from the CHARMM36 forcefield^133^ and assigned using PROPKA^134^ and PBD2PQR.^135^

#### Computing electrophoretic mobilities

To obtain the concentration-dependent ionic mobility 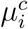 from fitted functions, it must first be derived from the salt’s molar conductivity Λ and the ion’s transport number *t*_*i*_ before it can be fitted^86^

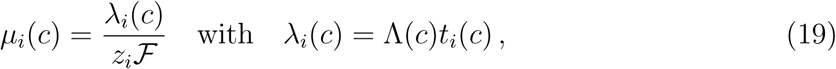

where *λ*_*i*_(*c*) is the specific molar conductivity of ion *i*.

#### Computing the simulated ionic current and electro-osmotic flow rate

The simulated ionic current *I*_sim_ at steady-state was computed by

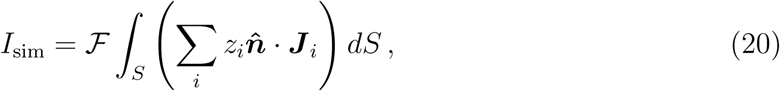

with *z*_*i*_ the charge number and ***J*** _*i*_ the total flux of each ion *i* across *cis* reservoir boundary *S*, ℱ the Faraday constant (96 485 C · mol^−1^) and 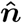 the unit vector normal to *S*. Similarly, the volumetric flow rate, *i.e.*, the volume of water passing through the pore per unit time, is given by

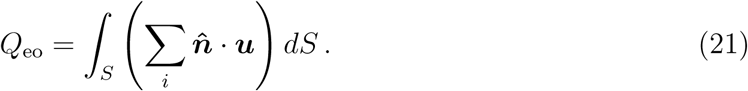

#### ClyA expression and purification

ClyA-AS monomers were expressed, purified and oligomerized using methods described in detail elsewhere.^25,96^ Briefly, *E. cloni* EXPRESS BL21 (DE3) cells (Lucigen Corporation, Middleton, USA) were transformed with a pT7 plasmid containing the ClyA-AS gene, followed by overexpression after induction with 1 mM isopropyl β-D-1-thiogalactopyranoside (Sigma-Aldrich, Zwijndrecht, The Netherlands). The ClyA monomers were purified using Ni^+^-NTA affinity chromatography and oligomerized by incubation in 0.2 % D-maltoside n-dodecyl-β-D-maltopyranoside (Sigma-Aldrich, Zwijndrecht, The Netherlands) for 30 minutes at 37°C. Pure ClyA-AS type-I (12-mer) nanopores were obtained using native PAGE on a 4-15% gradient gel (Bio-Rad, Veenendaal, The Netherlands) and subsequent excision of the correct oligomer band.

#### Recording of single-channel current-voltage curves

Experimental current-voltage curves where measured using single-channel electrophysiology, as detailed elsewhere. ^25,35,96^ Briefly, a black lipid bilayer was inside a ≈100 µm diameter aperture in a thin teflon film separating two buffered electrolyte compartments by painting with 1,2-diphytanoyl-snglycero-3-phosphocholine (DPhPC, Avanti Polar Lipids, Alabaster, USA). Minute amounts (≈) of the purified ClyA-AS type I oligimer were then added to the grounded *cis* reservoir and allowed to insert into the lipid bilayer. Single-channel current-voltage curves were recorded using a custom pulse protocol of the Clampex 10.4 software package connected to AxoPatch 200B patch-clamp amplifier via a Digidata 1440A digitizer (all from Molecular Devices, San Jose, USA). Data was acquired at 10 kHz and filtered using a 2-kHz low bandpass filter. Measurements at different ionic strengths were performed at ≈ 25°C in aqueous NaCl solutions, buffered at pH 7.5 using 10 mM MOPS (Sigma-Aldrich, Zwijndrecht, The Netherlands).

## Acknowledgement

K.W., J.H. and P.V.D. acknowledge the financial support by the FWO (grant number 3E130054). G.M. has received funding from the European Research Council (ERC) under the European Union’s Horizon 2020 research and innovation programme (Grant agreement No. 726151). The authors thank Niels Verellen and Chang Chen for their valuable feedback during discussions.

## Supporting Information Available

The supplementary info contains the Extended materials and methods, with details on the fitting of the electrolyte properties and the calculation of the pore averaged values, and the weak forms of the ePNP-NS equations. It also contains additional results, including a figures about ionic current rectification and the electro-osmotic pressure distribution inside the pore, and a table detailing the peak values of the radial electrostatic potential inside ClyA.

## Graphical TOC Entry

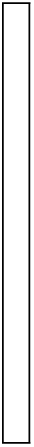

