## Supplementary text and figures for "Modeling of Ion and Water Transport in the Biological Nanopore ClyA"

‡*imec, Kapeldreef 75, B-3001 Leuven, Belgium*

¶*KU Leuven, Department of Physics and Astronomy, Celestijnenlaan 200D, B-3001 Leuven, Belgium*

§*University of Groningen, Groningen Biomolecular Sciences & Biotechnology Institute, 9747 AG, Groningen, The Netherlands*

### Contents

**1 Extended materials and methods** **S-3**

1.1 ClyA-AS mutations w.r.t. the wild-type . . . . . . . . . . . . . . . . . . . . S-3

1.2 Fitting of the electrolyte properties . . . . . . . . . . . . . . . . . . . . . . . S-3

1.3 Surface integration to compute pore averaged values . . . . . . . . . . . . . . S-5

1.4 Weak forms of the ePNP-NS equations . . . . . . . . . . . . . . . . . . . . . S-7

1.5 Estimating the bulk conductance of ClyA . . . . . . . . . . . . . . . . . . . . S-10

**2 Extended results** **S-12**

2.1 Ionic current rectification . . . . . . . . . . . . . . . . . . . . . . . . . . . . S-12

2.2 Peak values of the radial potential profiles inside ClyA . . . . . . . . . . . . S-12

2.3 Pressure distribution inside ClyA . . . . . . . . . . . . . . . . . . . . . . . . S-13

**References** **S-14**

### 1 Extended materials and methods

#### 1.1 ClyA-AS mutations w.r.t. the wild-type

**Table S1.** Mutations of the ClyA-AS variant compared to the *S. tyhpii* wild-type.

| Position | WT | AS |
|---|---|---|
| 8 | Lys | Gln |
| 15 | Asn | Ser |
| 38 | Gln | Lys |
| 57 | Ala | Glu |
| 67 | Thr | Val |
| 87 | Cys | Ala |
| 90 | Ala | Val |
| 95 | Ala | Ser |
| 99 | Leu | Gln |
| 103 | Glu | Gly |
| 118 | Lys | Arg |
| 119 | Leu | Ile |
| 124 | Ile | Val |
| 125 | Thr | Lys |
| 136 | Val | Thr |
| 166 | Phe | Tyr |
| 172 | Lys | Arg |
| 185 | Val | Ile |
| 212 | Lys | Asn |
| 214 | Lys | Arg |
| 217 | Ser | Thr |
| 224 | Thr | Ser |
| 227 | Asn | Ala |
| 244 | Thr | Ala |
| 276 | Glu | Gly |
| 285 | Cys | Ser |
| 290 | Lys | Gln |

#### 1.2 Fitting of the electrolyte properties

The parameters of the fitting functions used to interpolate the experimental ion diffusion coefficients, mobilities and transport numbers, and the electrolyte viscosity, density and relative permittivity are given in Tab. S2 and the resulting curves are plotted in Fig. S1.

Note that since most of these functions are merely empirical fits with no physical meaning, and that they are solely used to interpolate and represent the experimental data.

Finally, for the concentration dependence of the relative permittivity we made use of the model proposed by Gavish et al.[4]

$$\varepsilon_r(\bar{c}) = \varepsilon_{r,0} - (\varepsilon_{r,0} - \varepsilon_{r,ms})\, L\left(\frac{3\alpha}{\varepsilon_{r,0} - \varepsilon_{r,ms}}\bar{c}\right) \tag{S1}$$

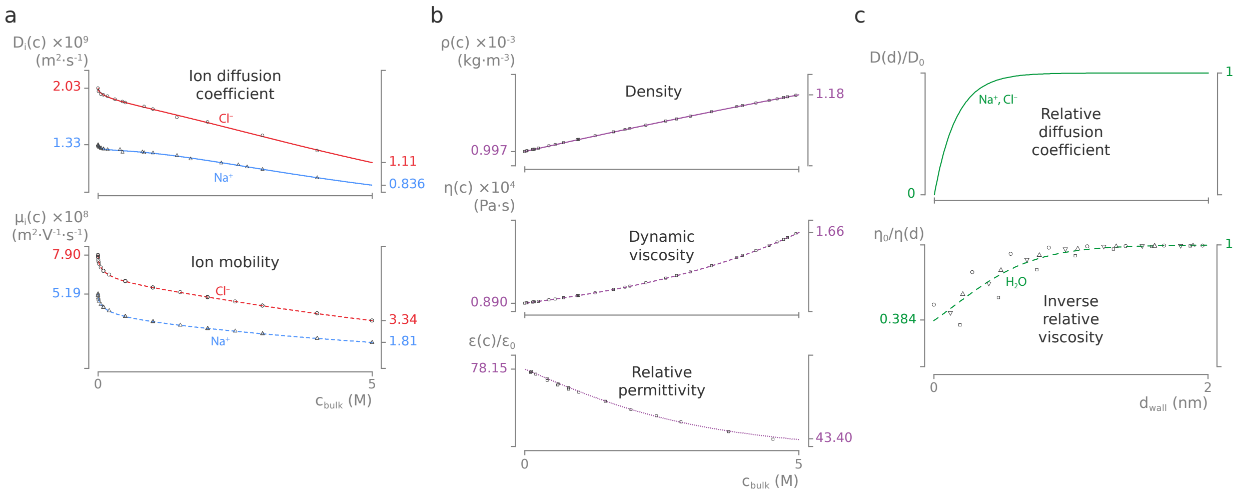


**Figure S1. Concentration and positional dependent electrolyte properties.** (a) Dependency of the ion self-diffusion coefficients (top, $Na^+$: blue solid line and $Cl^-$: red solid line) and the ion electrophoretic mobilities (bottom, $Na^+$: blue dashed line and $Cl^-$: red dashed line) on the bulk NaCl concentration. Lines represent empirical fits to experimental literature data (markers). The number next to the left and right axes correspond to the values at infinite dilution (0 M) and saturation (≈5 M), respectively. (b) Dependency of electrolyte density (top, purple solid line), viscosity (middle, purple dashed line) and relative permittivity (bottom, purple dotted line) on the bulk NaCl concentration. Lines represent empirical fits to experimental literature data (markers). (c) Dependency of the relative ion diffusion coefficient and the mobility (top, green solid line) and the relative viscosity (bottom, green dashed line) on the distance from the nanopore wall. The relative diffusion coefficient (and its mobility) declines sharply when an ion approaches within ≈1.0 nm of the protein wall. For water molecules, this phenomenon is modelled as a sharp increase of the fluid viscosity within ≈1.0 nm distance from the wall. The values for the diffusion coefficient were taken directly from ref. 1, who used an empirical fit on molecular dynamics data from ref. 2. The inverse relative viscosity data was taken directly from the molecular dynamics study in ref. 3, and fitted with a logistic function after offsetting for the protein hydrodynamic radius.

**Table S2.** Overview of the NaCl fitting parameters used for interpolation.

| | Fitting parameters | | | | | | |
|---|---|---|---|---|---|---|---|
| Property | $P_0$ | $P_1$ | $P_2$ | $P_3$ | $P_4$ | $R^2$ | References |
| $\mathcal{D}_+(c)$ | 1.334 | $2.02\pm0.14\times10^{-1}$ | $-3.05\pm0.41\times10^{-1}$ | $2.19\pm0.38\times10^{-1}$ | $-3.13\pm1.08\times10^{-2}$ | >0.99 | 5 |
| $\mathcal{D}_-(c)$ | 2.032 | $1.49\pm0.30\times10^{-1}$ | $-4.94\pm9.04\times10^{-2}$ | $3.40\pm8.26\times10^{-2}$ | $1.43\pm2.30\times10^{-2}$ | >0.99 | 5 |
| $\mu_+(c)$ | 5.192 | $7.91\pm0.06\times10^{-1}$ | $-3.53\pm0.17\times10^{-1}$ | $1.46\pm0.15\times10^{-1}$ | $9.23\pm3.89\times10^{-3}$ | >0.99 | 6–9 |
| $\mu_-(c)$ | 7.909 | $6.29\pm0.06\times10^{-1}$ | $-4.29\pm0.17\times10^{-1}$ | $2.12\pm0.14\times10^{-1}$ | $-1.07\pm0.37\times10^{-2}$ | >0.99 | 6–9 |
| $t_+(c)$ | 0.3963 | $9.38\pm1.64\times10^{-2}$ | $2.86\pm3.24\times10^{-3}$ | $-1.88\pm6.52\times10^{-2}$ | $4.51\pm2.75\times10^{-3}$ | 0.98 | 7,9–12 |
| $\eta(c)$ | 0.8904 | $7.56\pm0.27\times10^{-3}$ | $7.77\pm0.04\times10^{-2}$ | $1.19\pm0.01\times10^{-2}$ | $5.95\pm0.35\times10^{-4}$ | >0.99 | 13 |
| $\varrho(c)$ | 0.997 | $4.06\pm0.01\times10^{-2}$ | $-6.39\pm0.16\times10^{-4}$ | | | >0.99 | 13 |
| $\varepsilon(c)$ | 78.15 | $3.08\pm0.00\times10^{1}$ | $1.15\pm0.00\times10^{1}$ | | | | 4 |
| $\mathcal{D}_+(d)$ | | 6.2 | 0.01 | | | | 1,2,14 |
| $\mathcal{D}_-(d)$ | | 6.2 | 0.01 | | | | 1,2,14 |
| $\mu_+(d)$ | | 6.2 | 0.01 | | | | 1,2,14 |
| $\mu_-(d)$ | | 6.2 | 0.01 | | | | 1,2,14 |
| $\eta(d)$ | | $3.36\pm0.23$ | $1.47\pm0.23\times10^{-1}$ | | | 0.97 | 3 |

#### 1.3 Surface integration to compute pore averaged values

The average pore values for quantity of interest $X$ was computed by

$$\langle X \rangle_\alpha = \frac{\iint_{V_\alpha} \beta_\alpha X \, dr \, dz}{\iint_{V_\alpha} \beta_\alpha \, dr \, dz} \,, \tag{S2}$$

where

$$\alpha = \begin{cases} \text{p}, & d \geq 0 \text{ nm, average over the entire pore} \\ \text{b}, & d > 0.5 \text{ nm, average over the pore `bulk'} \\ \text{s}, & d \leq 0.5 \text{ nm, average over the pore `surface'} \end{cases} \tag{S3}$$

and

$$\beta_{\mathrm{p}} = \begin{cases} 1, & \text{if } -1.85 \leq z \leq 12.25 \text{ and } r \leq r_{\mathrm{p}}(z) \\ 0, & \text{otherwise} \end{cases} \tag{S4}$$

$$\beta_{\mathrm{b}} = \begin{cases} 1, & \text{if } -1.85 \leq z \leq 12.25 \text{ and } r \leq r_{\mathrm{p}}(z) \text{ and } d > 0.5 \\ 0, & \text{otherwise} \end{cases} \tag{S5}$$

$$\beta_{\mathrm{s}} = \begin{cases} 1, & \text{if } -1.85 \leq z \leq 12.25 \text{ and } r \leq r_{\mathrm{p}}(z) \text{ and } d \leq 0.5 \\ 0, & \text{otherwise} \end{cases} \tag{S6}$$

with $d$ the distance from the nanopore wall and $r_{\mathrm{p}}(z)$ is the radius of the pore at height $z$.

#### 1.4 Weak forms of the ePNP-NS equations

To solve partial differential equations with the finite element method, we must be derive their weak form. This is achieved through multiplication of the equation with an arbitrary test function and integration over their relevant domains and boundaries (Fig. S2). The full computational domain of our model ($\Omega$) is subdivided into domains for the pore ($\Omega_p$), the lipid bilayer ($\Omega_m$) and the electrolyte reservoir ($\Omega_w$). The relevant boundaries of these domains are also indicated, i.e. the reservoir's exterior edges at the *cis* $\Gamma_{w,c}$) and *trans* $\Gamma_{w,t}$) sides, the outer edge of the lipid bilayer ($\Gamma_m$) and the interface of the fluid with the nanopore and the bilayer ($\Gamma_{p+m}$).

The following paragraphs detail the equations that were used in this paper, together with their corresponding weak forms.

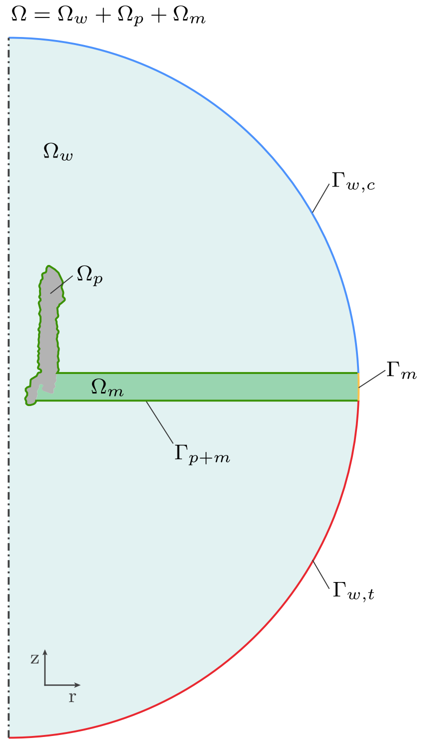


**Figure S2. Computational domains and boundaries.** The full computational domain ($\Omega$) of the model is subdivided into various subdomains ($\Omega_x$) and their limiting boundaries ($\Gamma_x$).

**Poisson equation.** The global potential distribution is described by the Poisson equation (PE)[15]

$$\nabla \cdot \boldsymbol{D} = -\left(\rho^f_{\text{pore}} + \rho_{\text{ion}}\right) \quad \text{with } \boldsymbol{D} = \varepsilon_0 \varepsilon_r \nabla\varphi \ , \tag{S7}$$

with $\boldsymbol{D}$ the electrical displacement field, $\varphi$ the electric potential, $\varepsilon_0$ the vacuum permittivity ($8.85419 \times 10^{-12}$ F m$^{-1}$), and $\varepsilon_r$ the local relative permittivity of the medium. $\rho^f_{\text{pore}}$ and $\rho_{\text{ion}}$ are the fixed (due to the pore) and mobile (due to the ions) charge distributions, respectively.

Multiplication of Eq. (S7) with the potential test function $\psi$ and integration over the entire model $\Omega = \Omega_w + \Omega_p + \Omega_m$ gives

$$\int_\Omega \left[\nabla \cdot \boldsymbol{D}\right] \psi \, d\Omega = -\int_\Omega \left[\rho^f_{\text{pore}} + \rho_{\text{ion}}\right] \psi \, d\Omega \ , \tag{S8}$$

which, after applying the Gauss divergence theorem, yields the final weak formulation

$$\int_\Omega \left[\nabla\psi \cdot \boldsymbol{D}\right] d\Omega - \int_{\Gamma_{\text{PE}}} \left[\psi \boldsymbol{D} \cdot \boldsymbol{n}\right] d\Gamma_{\text{PE}} = \int_\Omega \left[\psi \rho^f_{\text{pore}}\right] d\Omega + \int_\Omega \left[\psi \rho_{\text{ion}}\right] d\Omega \ , \tag{S9}$$

with boundaries $\Gamma_{\text{PE}} = \Gamma_{w,c} + \Gamma_{w,t} + \Gamma_m$ and $\boldsymbol{n}$ their normal vector. The boundary integrals at $\Gamma_{w,c}$ and $\Gamma_{w,t}$ are evaluated using the Dirichlet boundary conditions (BCs) $\varphi = 0$ and $\varphi = V_{\text{b}}$, respectively. A zero charge BC, $\boldsymbol{n} \cdot \boldsymbol{D} = 0$, is used for the integral at $\Gamma_m$.

**Size-modified Nernst-Planck equation.** The total ionic flux $\boldsymbol{J}_i$ of ion $i$ at steady-state is expressed by the size-modified Nernst-Planck equation (smNPE)[15]

$$\frac{\partial c_i}{\partial t} = -\nabla \cdot \boldsymbol{J}_i = -\nabla \cdot \left(\mathcal{D}_i \nabla c_i + z_i \mu_i c_i \nabla\varphi + \boldsymbol{\beta_i} c_i - \boldsymbol{u} c_i\right) \quad \text{with } \boldsymbol{\beta_i} = \frac{a_i^3/a_0^3 \sum\limits_j a_j^3 \nabla c_j}{1 - \sum\limits_j N_{\text{A}} a_j^3 c_j} \ , \tag{S10}$$

with ion diffusion coefficient $\mathcal{D}_i$, concentration $c_i$, charge number $z_i$, mobility $\mu_i$, electrostatic potential $\varphi$, steric saturation factor $\beta_i$ and fluid velocity $\boldsymbol{u}$. $N_{\text{A}}$ is Avogadro's constant

($6.022 \times 10^{23}$ mol$^{-1}$) and $a_i$ and $a_0$ are the limiting cubic diameters for ions and water, respectively. Using the ion concentration test function $d_i$, the weak form of Eq. (S10) becomes

$$
\begin{aligned}
\int_{\Omega_w} \left[\frac{\partial c_i}{\partial t}\right] d_i \, d\Omega_w &= \int_{\Omega_w} \left[-\nabla \cdot \boldsymbol{J}_i\right] d_i \, d\Omega_w \\
\int_{\Omega_w} \left[d_i \frac{\partial c_i}{\partial t}\right] d\Omega_w &= \int_{\Omega_w} \left[\nabla d_i \cdot \boldsymbol{J}_i\right] d\Omega_w - \int_{\Gamma_{\mathrm{NP}}} \left[d_i \boldsymbol{J}_i \cdot \boldsymbol{n}\right] d\Gamma_{\mathrm{NP}} \\
&= \int_{\Omega_w} \left[\nabla d_i \cdot \left(\mathcal{D}_i \nabla c_i + z_i \mu_i c_i \nabla \varphi + \boldsymbol{\beta_i} c_i - \boldsymbol{u} c_i\right)\right] d\Omega_w \\
&\quad - \int_{\Gamma_{\mathrm{NP}}} \left[d_i \left(\mathcal{D}_i \nabla c_i + z_i \mu_i c_i \nabla \varphi + \boldsymbol{\beta_i} c_i - \boldsymbol{u} c_i\right) \cdot \boldsymbol{n}\right] d\Gamma_{\mathrm{NP}} \ .
\end{aligned}
\tag{S11}
$$

The integrals on the boundaries $\Gamma_{\mathrm{NP}} = \Gamma_{w,c} + \Gamma_{w,t} + \Gamma_{p+m}$ are evaluated using the Dirichlet boundary condition $c_i = c_{\mathrm{s}}$ for $\Gamma_{w,c}$ and $\Gamma_{w,t}$, and the no flux boundary condition $\boldsymbol{n} \cdot \boldsymbol{J}_i = 0$ for $\Gamma_{p+m}$

**Variable density and viscosity Navier-Stokes equation.** The steady-state, laminar fluid flow of an incompressible fluid with a variable density and viscosity is given by the system of equations[16]

$$\boldsymbol{u} \cdot \nabla \varrho = 0 \tag{S12}$$

$$(\boldsymbol{u} \cdot \nabla)(\varrho \boldsymbol{u}) + \nabla \cdot \sigma_{ij} = \boldsymbol{F} \quad \text{with } \sigma_{ij} = p\mathbf{I} - \eta \left[\nabla \boldsymbol{u} + (\nabla \boldsymbol{u})^{\mathsf{T}}\right] \tag{S13}$$

$$\nabla \cdot (\varrho \boldsymbol{u}) - \boldsymbol{u} \cdot \nabla \varrho = 0 \ , \tag{S14}$$

with fluid velocity $\boldsymbol{u}$, density $\varrho$, hydrodynamic stress tensor $\sigma_{ij}$, viscosity $\eta$, pressure $p$ and body force $\boldsymbol{F}$. The pressure test function $q$ is used to derive the weak forms of Eq. (S12)

$$\int_{\Omega_w} \left[\boldsymbol{u} \cdot \nabla \varrho\right] q \, d\Omega_w = \int_{\Omega_w} \left[q \boldsymbol{u} \cdot \nabla \varrho\right] d\Omega_w = 0 \ , \tag{S15}$$

and Eq. (S14)

$$\begin{aligned}\int_{\Omega_w} [\nabla \cdot (\varrho \boldsymbol{u}) - \boldsymbol{u} \cdot \nabla \varrho]\, q \, d\Omega_w &= \int_{\Omega_w} [\nabla \cdot (\varrho \boldsymbol{u})]\, q \, d\Omega_w - \int_{\Omega_w} [q \boldsymbol{u} \cdot \nabla \varrho] \; d\Omega_w \\ &= \int_{\Omega_w} [\nabla q \cdot (\varrho \boldsymbol{u})] \; d\Omega_w - \int_{\Gamma_{\mathrm{NS}}} [q\, (\varrho \boldsymbol{u}) \cdot \boldsymbol{n}] \; d\Gamma_{\mathrm{NS}} - \int_{\Omega_w} [q \boldsymbol{u} \cdot \nabla \varrho] \; d\Omega_w \ ,\end{aligned} \tag{S16}$$

while for Eq. (S13) we use the velocity test function $\boldsymbol{v} = [v_r, v_\phi, v_z]$

$$\begin{aligned}\int_{\Omega_w} [(\boldsymbol{u} \cdot \nabla)\, (\varrho \boldsymbol{u}) + \nabla \cdot \sigma_{ij}] \cdot \boldsymbol{v} \, d\Omega_w = \int_{\Omega_w} \boldsymbol{F} \cdot \boldsymbol{v} \, d\Omega_w \\ \int_{\Omega_w} [(\boldsymbol{u} \cdot \nabla)\, (\varrho \boldsymbol{u}) \cdot \boldsymbol{v}] \; d\Omega_w - \int_{\Omega_w} [\sigma_{ij} \cdot \nabla \boldsymbol{v}] \; d\Omega_w + \int_{\Gamma_{\mathrm{NS}}} [\boldsymbol{v} \cdot \sigma_{ij} \cdot \boldsymbol{n}] \; d\Gamma_{\mathrm{NS}} = \int_{\Omega_w} \boldsymbol{F} \cdot \boldsymbol{v} \, d\Omega_w \ ,\end{aligned} \tag{S17}$$

with boundaries $\Gamma_{\mathrm{NS}} = \Gamma_{w,c} + \Gamma_{w,t} + \Gamma_{p+m}$. The no-slip Dirichlet BC $\boldsymbol{u} = 0$ is applied to $\Gamma_{p+m}$, and the no normal stress $\sigma_{ij} \boldsymbol{n} = 0$ is used for $\Gamma_{w,c}$ and $\Gamma_{w,t}$.

#### 1.5 Estimating the bulk conductance of ClyA

The ionic current flowing through a nanopore can be computed using Ohm's law

$$I(c_{\mathrm{s}}, V_{\mathrm{b}}) = G_{\mathrm{bulk}}(c_{\mathrm{s}}) V_{\mathrm{b}} \tag{S18}$$

with $G_{\mathrm{bulk}}$ the nanopore conductance. A naive estimate of $G_{\mathrm{bulk}}$ can be obtained by modelling it as a fluidic resistor, represented by a fluid-filled cylinder.[17] Because ClyA has an asymmetric shape, it is best approximated by a two resistors in series: a tall, wide cylinder (*cis lumen*) on top of a narrow short cylinder (*trans* constriction).[18] Hence,

$$G_{\mathrm{bulk}(c_{\mathrm{s}})} = \frac{\sigma}{4\pi}(c_{\mathrm{s}}) \left( \frac{l_{\mathrm{cis}}}{d_{\mathrm{cis}}^2} + \frac{l_{\mathrm{trans}}}{d_{\mathrm{trans}}^2} \right) \ , \tag{S19}$$

with $\sigma$ the (concentration dependent) bulk electrolyte conductivity. The *cis* and *trans* chambers have have heights and diameters of $l_{\text{cis}} = 10\,\text{nm}$ and $l_{\text{trans}} = 4\,\text{nm}$, and $d_{\text{cis}} = 10\,\text{nm}$ and $d_{\text{trans}} = 4\,\text{nm}$, respectively.[18]

### 2 Extended results

#### 2.1 Ionic current rectification

The ionic current rectification (ICR, $\alpha$, Fig. S3) represents the relative conductivity of the nanopore at identical, but opposing bias voltages

$$\alpha(V_b) = \frac{G(+V_b)}{G(-V_b)} \text{ , with } V_b \geq 0 \text{ .}$$

Hence, for $\alpha > 0$, the current is higher at positive bias voltages, and vice versa for $\alpha < 0$. The ICR phenomenon results from an asymmetry in the ionic conductance pathways (*e.g.*, moving from *cis* to *trans* is not equivalent as moving from *trans* to *cis*). In turn, this stems from the intrinsic asymmetries of the nanopore itself (*i.e.*, geometry and charge distribution) or its response to the electric field (*i.e.*, gating or electrostriction).

The heatmap of the simulated (ePNP-NS) $\alpha$ (Fig. S3a) reveals that $\alpha > 0$ between 0 mV to 200 mV and 0.005 M to 5 M. It increases monotonically with increasing bias voltage and exhibits a maximum as a function of salt concentration at ≈0.15 M. The concentration dependency of $\alpha$ at 50 mV, 100 mV and 150 mV (Fig. S3b) shows that it increases monotonically with $c_s$ until it reaches a peak value, after which it decreases to approach unity ($\alpha = 1$) at saturating salt concentrations ($c_s \approx 5$ M). Even though the values produced by the ePNP-NS simulations do not fully match quantitatively to the experimental results, at least for $c_s < 0.5$ M, they do 1) reproduce the observed trends qualitatively and 2) exhibit much smaller errors relative to the PNP-NS results (data not shown).

#### 2.2 Peak values of the radial potential profiles inside ClyA

The peak values of the radial electrostatic potential $\langle\varphi\rangle_{\mathrm{rad}}$ at the *cis* entry, middle of the lumen and the *trans* constriction for 0.005, 0.05, 0.15, 0.5 and 5 M NaCl are summarized in Tab. S3.

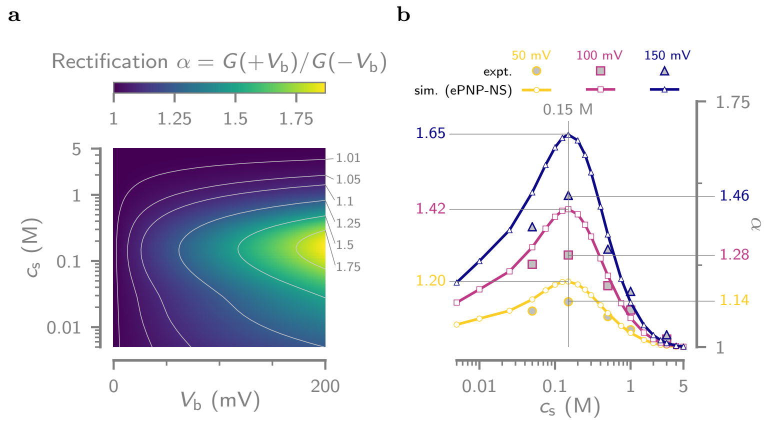


**Figure S3. Measured and simulated ionic rectificationof single ClyA nanopores.** (a) Contourplot of the ionic current rectification $\alpha = G(+V_b)/G(-V_b)$, computed from the ionic conductances in the ePNP-NS simulation, as a function of $V_b$ and $c_s$. (b) Comparision between the simulated (ePNP-NS) and measured (expt.) $\alpha$ as a function of $c_s$ for 50 mV, 100 mV and 150 mV.

**Table S3.** Peak radial potential.

| | $\langle\varphi\rangle_{rad}$ (mV) | | |
|---|---|---|---|
| | *cis* | lumen | *trans* |
| $c_s$ (M) | $z \approx 10$ nm | $z \approx 5$ nm | $z \approx 0$ nm |
| 0.005 | −80 | −108 | −144 |
| 0.05 | −34 | −50 | −86 |
| 0.15 | −19 | −29 | −57 |
| 0.5 | −9.3 | −14 | −30 |
| 5 | −1.9 | −1.7 | −4.2 |

#### 2.3 Pressure distribution inside ClyA

The electro-osmotic pressure distribution inside ClyA, as a consequence of the strong local enhancement of the ion concentration, is given in Fig. S4.

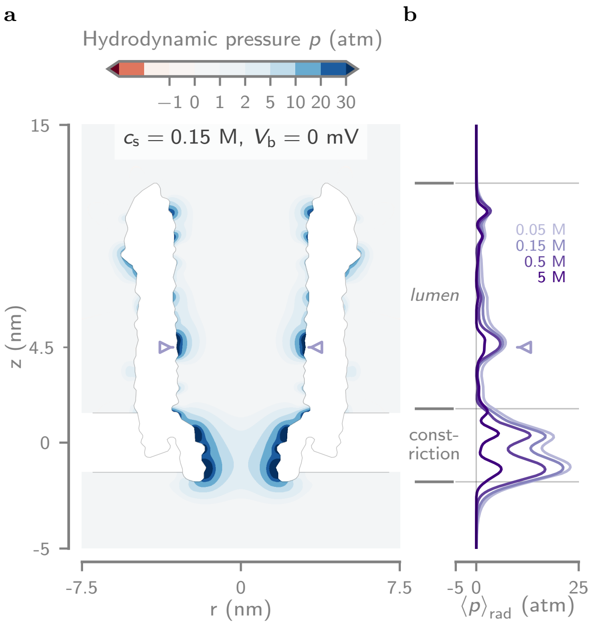


**Figure S4. Pressure distribution inside ClyA.** (a) Contourmap of the pressure at $c_s = 0.15\,\mathrm{M}$ and $V_b = 0\,\mathrm{mV}$, showing that the $Na^+$ concentration 'hotspots' near the pore wall result in build-up of electro-osmotic pressure (5 ATM to 30 ATM) inside the confined fluid. Pressure drops of such magnitude over the course of a few nanometers could potentially exert a significant force on a captured protein.[19] (b) The axial pressure profile and averaged along the the entire radius of the pore at $V_b = 0\,\mathrm{mV}$.
